# Brain Endothelial Cell TRPA1 Channels Initiate Neurovascular Coupling

**DOI:** 10.1101/2020.09.14.295600

**Authors:** Pratish Thakore, Michael G. Alvarado, Sher Ali, Amreen Mughal, Paulo W. Pires, Evan Yamasaki, Harry A. T. Pritchard, Brant E. Isakson, Cam Ha T. Tran, Scott Earley

**Affiliations:** Department of Pharmacology, Center for Molecular and Cellular Signaling in the Cardiovascular System, University of Nevada, Reno School of Medicine, Reno, NV 8957-0318, USA; Department of Pharmacology, College of Medicine, University of Vermont, Burlington, VT 05405, USA; Department of Physiology, College of Medicine, University of Arizona, Tucson, AZ 85724, USA; Institute of Cardiovascular Sciences, University of Manchester, M13 9PL Manchester, United Kingdom; Department of Molecular Physiology and Biological Physics, University of Virginia, Charlottesville, VA 22908, USA; Robert M. Berne Cardiovascular Research Center, University of Virginia, Charlottesville, VA 22908, USA; Department of Physiology & Cell Biology, Center for Molecular and Cellular Signaling in the Cardiovascular System, University of Nevada, Reno School of Medicine, Reno, NV 8957-0318, USA

**Keywords:** TRPA1 channels, Panx1 channels, purinergic signaling, conducted vasodilation, neurovascular coupling.

## Abstract

Blood flow regulation in the brain is dynamically regulated to meet the metabolic demands of active neuronal populations. Recent evidence has demonstrated that capillary endothelial cells are essential mediators of neurovascular coupling that sense neuronal activity and generate a retrograde, propagating, hyperpolarizing signal that dilates upstream arterioles. Here, we tested the hypothesis that transient receptor potential ankyrin 1 (TRPA1) channels in capillary endothelial cells are significant contributors to functional hyperemic responses that underlie neurovascular coupling in the brain. Using an integrative *ex vivo* and *in vivo* approach, we demonstrate the functional presence of TRPA1 channels in brain capillary endothelial cells, and show that activation of these channels within the capillary bed, including the post-arteriole transitional region covered by ensheathing mural cells, initiates a retrograde signal that dilates upstream parenchymal arterioles. Notably, this signaling exhibits a unique biphasic mode of propagation that begins within the capillary network as a short-range, Ca^2+^ signal dependent on endothelial pannexin-1 channel/purinergic P_2_X receptor communication pathway and then is converted to a rapid, inward-rectifying K^+^ channel-mediated electrical signal in the post-arteriole transitional region that propagates upstream to parenchymal arterioles. Two-photon laser-scanning microscopy further demonstrated that conductive vasodilation occurs *in vivo*, and that TRPA1 is necessary for functional hyperemia within the somatosensory cortex of mice. Together, these data establish a role for endothelial TRPA1 channels as sensors of neuronal activity and show that they respond accordingly by initiating a vasodilatory response that redirects blood to regions of metabolic demand.

## Introduction

Neurons within the brain lack significant intrinsic energy reserves; thus, the metabolic substrates for energy generation must be delivered to active regions in real time. However, substantial increases in global blood flow to the brain are limited by constraints imposed by the enclosing skull and the finite amount of total blood available for regional distribution. The brain therefore relies on dynamic local distribution of the available blood supply to the sites of highest metabolic activity. This vital process is accomplished by a complex interconnected network of cerebral pial arteries at the surface, parenchymal arterioles that provide a conduit to the interior, and a vast capillary network. Communication between active neurons and the cerebral microvasculature regulates activity-dependent increases in blood perfusion (functional hyperemia) in the brain through processes collectively referred to as “neurovascular coupling” (NVC) (Iadecola, 2017). One conceptual model of NVC that has received considerable research attention postulates that glutamate released during neurotransmission binds to G_q/11_ protein-linked metabotropic glutamate receptors (mGluRs) on perisynaptic astrocytic processes to stimulate Ca^2+^-dependent signaling pathways that ultimately cause the release of vasodilator substances from astrocytic endfeet encircling parenchymal arterioles (Attwell et al., 2010; Filosa, Bonev, & Nelson, 2004; Filosa et al., 2006; Iadecola & Nedergaard, 2007). These factors are purported to act directly on smooth muscle cells in the walls of arterioles to cause vasodilation and locally increase blood flow. Several recent studies have challenged this model, including one reporting that mGluR expression in astrocytes is undetectable after postnatal week 3, implying that this mechanism does not function in adults (Sun et al., 2013). Other reports have suggested that NVC can occur in the absence of astrocytic Ca^2+^ signaling (Jego, Pacheco-Torres, Araque, & Canals, 2014; Nizar et al., 2013) and that astrocytic Ca^2+^ signals lag vasodilator responses recorded *in vivo* in awake, behaving mice (Tran, Peringod, & Gordon, 2018). Longden et al. put forward a new paradigm for NVC, demonstrating that activation of inward-rectifying K^+^ (K_ir_) channels on brain capillary endothelial cells by extracellular K^+^ ions, released during neuronal activity, generates propagating electrical signals that cause dilation of upstream parenchymal arterioles (Longden et al., 2017). Given the essential nature of the NVC response for healthy brain function and life itself, it is reasonable to presume that multiple sensory modalities with overlapping and complementary roles operate within brain capillary endothelial cells to ensure the fidelity of neuronally driven functional hyperemia.

Here, we tested the hypothesis that transient receptor potential ankyrin 1 (TRPA1) channels in brain capillary endothelial cells contribute to functional hyperemia in the brain. Members of the TRP superfamily of cation channels act as molecular sensors of diverse physical and chemical stimuli (Flockerzi & Nilius, 2014). In cerebral pial arteries and parenchymal arterioles, activation of TRPA1 channels by endogenously produced lipid peroxide products, including 4-hydroxynonenal (4-HNE), leads to endothelium-dependent vasodilation (Earley, Gonzales, & Crnich, 2009; Pires & Earley, 2018; Qian, Francis, Solodushko, Earley, & Taylor, 2013; Sullivan et al., 2015; Trevisani et al., 2007). Using a combination of *ex vivo* and *in vivo* approaches, we find that activation of endothelial cell TRPA1 channels within capillaries, including post-arteriole transitional segments covered by enwrapping mural cells, generates retrograde propagating signals that lead to the dilation of upstream parenchymal arterioles. Specifically, we demonstrate that Ca^2+^ signals initiated by TRPA1 channels slowly propagate through the capillary network by a previously unknown pathway that is dependent on ionotropic purinergic P_2_X receptor activity and pannexin-1 (Panx1) expression. Within the post-arteriole region, slowly-propagating Ca^2+^ signals are converted to rapidly propagating electrical signals that cause dilation of upstream arterioles. Our data also show that endothelial cell TRPA1 channels contribute to blood flow regulation and functional hyperemia in the somatosensory cortex *in vivo*. We conclude that, under physiological conditions, TRPA1 channels in the brain endothelium are capable of mediating NVC.

## Results

### Functional TRPA1 channels are present in native brain capillary endothelial cells

Native cerebral capillary endothelial cells were freshly isolated using combined mechanical and enzymatic dispersion (Longden et al., 2017) and were patch-clamped in the conventional whole-cell configuration. Increasing extracellular [K^+^] to 60 mM stimulated Ba^2+^-sensitive K_ir_ currents in these cells, whereas NS309, a selective activator of small- and intermediate-conductance Ca^2+^-activated K^+^ channels (SK and IK, respectively) had no effect (Supplemental Figure 1A to D). These attributes are consistent with those reported for native brain capillary endothelial cells in a prior study (Longden et al., 2017).

Non-selective cation currents in native brain capillary endothelial cells were recorded by patch-clamp electrophysiology in the whole-cell configuration using symmetrical cation gradients established by intracellular and extracellular solutions. In isolated cells, application of voltage ramps from −80 to +80 mV over 300 ms in the presence of the endogenous TRPA1 agonist 4-HNE (100 nM) elicited dually rectifying currents with a reversal potential near 0 mV (Figure 1A and B). The selective TRPA1 inhibitor HC-030031 (10 µM) significantly attenuated this current. To confirm that 4-HNE actions are specific for TRPA1, we generated endothelial cell-specific TRPA1-knockout (*Trpa1-*ecKO) mice by crossing mice homozygous for *loxP* sites flanking the region within *Trpa1* encoding the S5 and S6 transmembrane domains (*Trpa1^fl/fl^*) with mice heterozygous for TEK tyrosine kinase (*Tek*) promoter-driven expression of Cre recombinase (*Tek^cre^*). Our prior study demonstrated that TRPA1 expression is undetectable by Western blotting in cerebral arteries from *Trpa1-*ecKO mice (Sullivan et al., 2015). *Trpa1^fl/fl^* mice, homozygous for floxed *Trpa1* but without expression of Cre recombinase, were used as controls. We found that 4-HNE did not stimulate cation currents in brain capillary endothelial cells isolated from *Trpa1*-ecKO mice (Figure 1C to F), supporting the selectivity of 4-HNE and the expression of functional TRPA1 channels in native brain capillary endothelial cells from wild-type and *Trpa1^fl/fl^* mice. Control patch-clamp experiments in which extracellular [K^+^] was raised to 60 mM to evoke K_ir_ currents (Longden et al., 2017) further confirmed the viability of capillary endothelial cells isolated from *Trpa1*-ecKO mice (Figure 1C to F).

**Figure 1.**
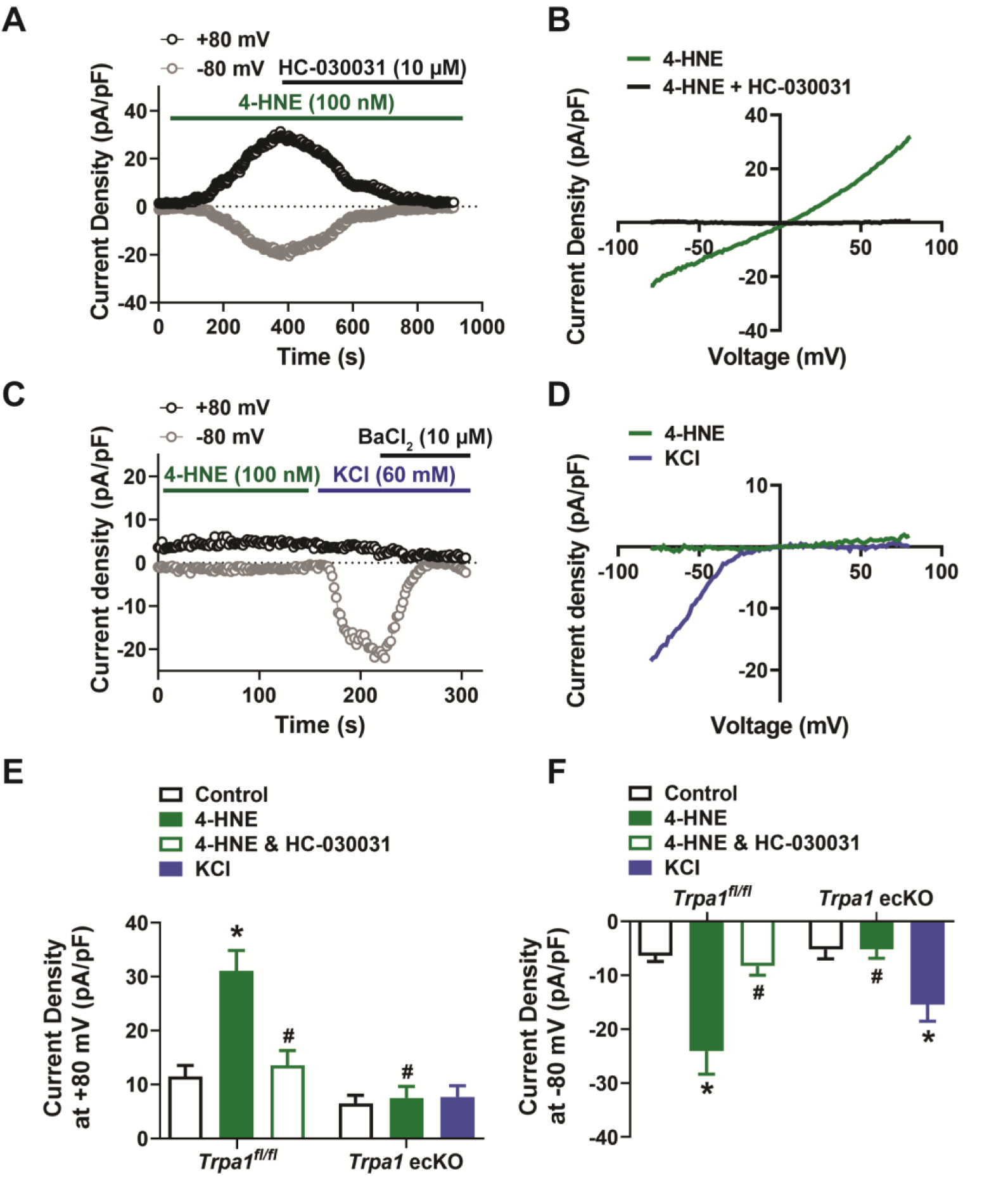
TRPA1 channels are functionally expressed in native brain capillary endothelial cells. A and B) Representative current versus time trace (A) and *I-V* relationship (B) from a whole-cell patch-clamp electrophysiology experiment demonstrating that the TRPA1 activator 4-HNE (100 nM) elicited a current in a native capillary endothelial cell isolated from a control *Trpa1^fl/fl^* mouse that was blocked by the selective TRPA1 antagonist HC-030031 (10 µM). C and D) Representative current versus time trace (C) and *I-V* relationship (D) demonstrating that 4-HNE was unable to elicit a current in a native capillary endothelial cell from a *Trpa1*-ecKO mouse. Cell viability was confirmed by evoking K_ir_ currents with raised extracellular KCl (60 mM) that were sensitive to BaCl_2_ (10 µM). E and F) Summary data showing that the TRPA1 current produced by 4-HNE at +80 mV (E) and −80 mV (F) was blocked by HC-030031 and was absent in cells from *Trpa1*-ecKO mice (*Trpa1^fl/fl^*, n = 12 cells from 5 animals; *Trpa1*-ecKO, n = 8 cells from 3 animals; *P < 0.05).

### Stimulation of TRPA1 channels in brain capillary endothelial cells induces dilation of upstream parenchymal arterioles

To determine if activation of capillary endothelial cell TRPA1 channels relaxes the upstream vasculature, we utilized a recently described *ex vivo* cerebral microvascular preparation in which microvascular segments with intact capillary branches are isolated from the brain, cannulated with micropipettes, and pressurized to 40 mmHg (Longden et al., 2017) (Figure 2A). In these experiments, a micropipette attached to a picospritzer was placed adjacent to the capillary extremities, which were stimulated by locally applying pulses (7 seconds; pressure, ∼10 psi) of various substances dissolved in artificial cerebral spinal fluid (aCSF). The spread of the picospritzed solution was assessed by pulsing aCSF containing 1% (w/v) Evans Blue dye. Control experiments showed that dye ejected from the micropipette was localized to the capillary region and did not spread to the upstream parenchymal arteriole segment (Supplemental Movie 1). To validate the viability of this preparation, we focally applied pulses of aCSF containing elevated [K^+^] (10 mM) onto capillary extremities. This maneuver induced a transient, reversible, and reproducible dilation of the upstream arteriole segment, as has been shown previously (Longden et al., 2017). This response was attenuated by incubating the *ex vivo* preparation with BaCl_2_ (30 µM) to inhibit K_ir_ channels (Supplemental Figure 2A and B). Picospritzing capillary extremities with standard aCSF (3 mM K^+^) did not affect the diameter of upstream arteriole segments (Supplemental Figure 2C and D). Additional control studies comparing the dilatory response of upstream arterioles to elevated [K^+^] application before and after severing capillaries from the upstream arteriole segment (Supplemental Figure 3A) showed that interruption of this connection prevented elevated [K^+^]-induced dilation of upstream arterioles (Supplemental Figure 3B). These data demonstrate the viability of our microvascular preparation and validate conducted vasodilation in response to stimulation of capillaries with elevated [K^+^], as previously reported by Longden et al. (Longden et al., 2017).

**Figure 2.**
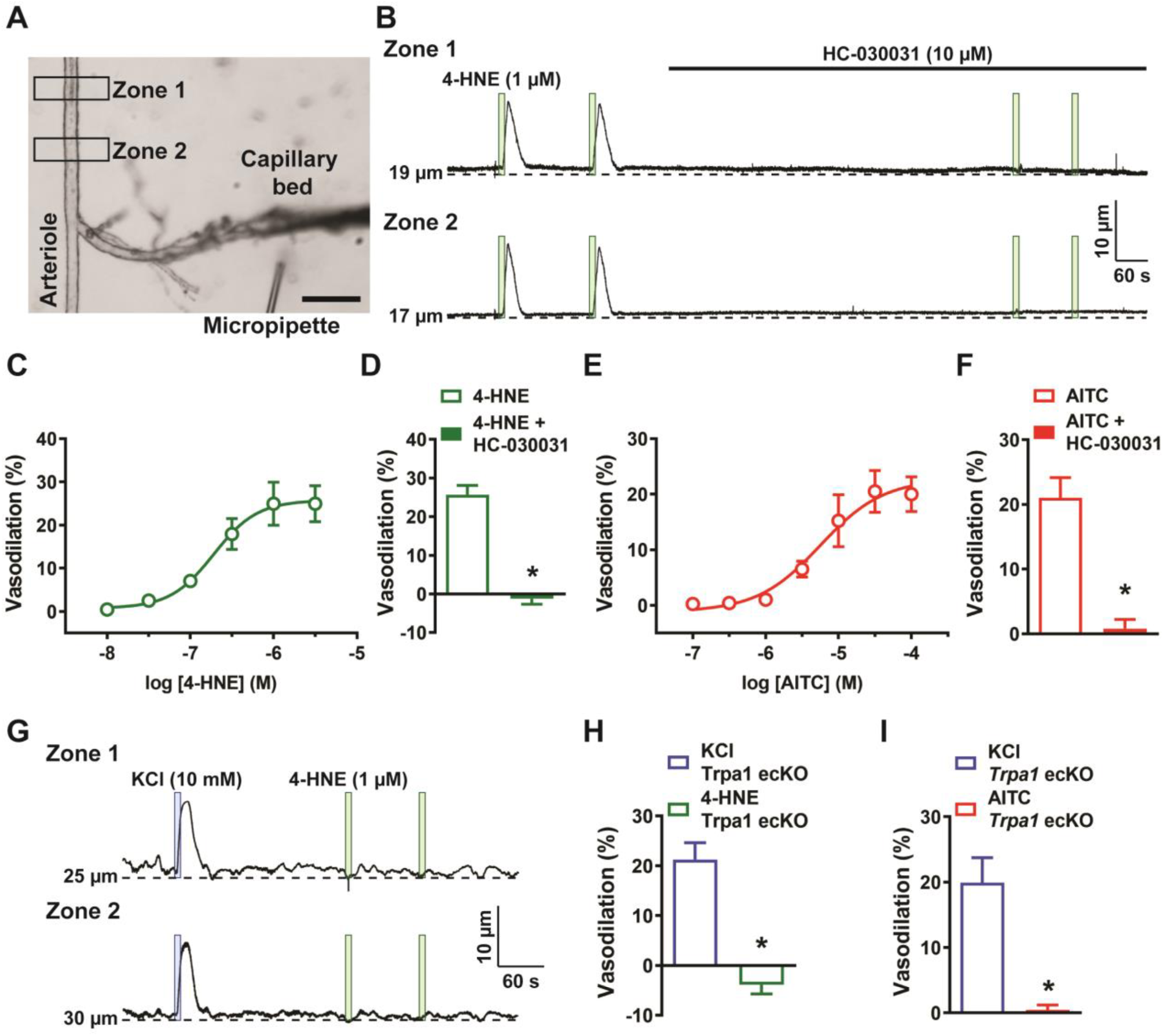
Capillary TRPA1 channels initiate conducted vasodilation in the cerebral microcirculation. A) Representative image of an *ex vivo* microvascular preparation consisting of an intact parenchymal arteriole with attached capillaries. Scale bar = 100 µm. Drugs were directly applied onto capillary extremities with a micropipette. B) Representative traces showing that application of 4-HNE (1 µM; green box) directly onto capillary extremities produced a reproducible increase in lumen diameter in two different areas (Zone 1 and Zone 2) on the upstream arteriole segment in microvascular preparations isolated from wild-type mice. The dilation produced by 4-HNE was blocked the selective TRPA1 antagonist HC-030031 (10 µM). C) Concentration-response curve produced by locally applying 4-HNE to capillary extremities over a concentration range of 10 nM to 30 µM (n = 6 preparations from 5 animals). An EC_50_ of 190 nM was calculated from the plot of a non-linear regression curve. D) Summary data showing that dilation to 4-HNE (1 µM) was blocked by HC-030031 (10 µM) (n = 9 preparations from 6 animals; *P < 0.05). E) Concentration-response curve produced by locally applying the TRPA1 agonist AITC to capillary extremities over a concentration range of 100 nM to 100 µM. An EC_50_ of 6 µM was calculated from the plot of a non-linear regression curve (n = 6 preparations from 5 animals). F) Summary data showing that dilation to AITC (30 µM) was blocked by HC-030031 (10 µM) (n = 7 preparations from 5 animals; *P < 0.05). G) Representative traces showing that application 4-HNE (1 µM; green box) onto capillary extremities was unable to dilate the upstream arteriole in microvascular preparations from *Trpa1*-ecKO mice, whereas application of KCl (10 mM; blue box) effectively dilated the upstream arteriole in these preparations. H and I) Summary data showing that neither 4-HNE (1 µM) (n = 7 preparations from 5 animals; *P < 0.05) (H) nor AITC (30 µM) (n = 7 preparations from 4 animals; *P < 0.05) (I) evoked dilation of upstream arterioles in preparations from *Trpa1*-ecKO mice.

TRPA1 channels in capillary segments of the *ex vivo* microvascular preparation were stimulated by focally applying pulses of various TRPA1-activating compounds dissolved in aCSF. Application of 4-HNE onto capillary extremities dilated upstream arterioles in *ex vivo* preparations from wild-type mice (Figure 2B) in a concentration-dependent manner. The EC_50_ for 4-HNE-induced dilation was 190 nM, and the maximal response was achieved at 1 µM; thus, this latter concentration was used in subsequent investigations (Figure 2C). Dilation of parenchymal arterioles in response to capillary-applied 4-HNE was significantly attenuated by superfusing the *ex vivo* preparation with HC-030031 (Figure 2B and 2D). Application of the TRPA1-activating compound, allyl isothiocyanate (AITC), to capillary beds also produced a concentration-dependent dilation of upstream arterioles (Supplemental Movie 2). The EC_50_ of AITC-induced arteriole dilation was 6 µM, and the maximal response was achieved at a concentration of 30 µM (Figure 2E). HC-030031 also inhibited upstream dilation in response to AITC (30 µM) (Figure 2F). In addition, the dilatory response of upstream arterioles to stimulation of capillary extremities with AITC was absent after severing the connection between capillaries and the upstream arteriole (Supplemental Figure 3C), providing evidence that this response requires intercellular signal propagation. In *ex vivo* preparations obtained from *Trpa1*-ecKO mice, stimulation of capillaries with elevated [K^+^] evoked dilation of upstream arterioles (Figure 2G to I); however, neither 4-HNE (Figure 2G and H) nor AITC (Figure 2I) had any effect at concentrations that produced maximal dilation of microvascular preparations from control animals. Together, these data demonstrate that activation of TRPA1 channels in brain capillary endothelial cells produces a signal that propagates to upstream parenchymal arterioles to cause dilation.

### Biphasic propagation velocity of conducted vasodilator signals initiated by TRPA1

We next used a genetic fluorescence reporter strategy to image the cell types present in our *ex vivo* cerebral microvascular preparation. Mice heterozygous for a membrane bound tdTomato cassette (*mT*) express red fluorescence in all cell types (Muzumdar, Tasic, Miyamichi, Li, & Luo, 2007). This cassette is flanked by *loxP* sites such that, upon Cre-mediated recombination, the *mT* cassette is excised, allowing expression of a membrane-targeted EGFP cassette (*mG*) in *Cre*-expressing cells only (Muzumdar et al., 2007). Mice expressing *mT/mG* were crossed with mice heterozygous for vascular endothelial cadherin (VEC)-*cre*, resulting in EGFP expression exclusively in endothelial cells and expression of tdTomato in other cell types. Epifluorescence images obtained from these mice showed an interconnected network of endothelial cells encompassed by mural cells (Figure 3A). In agreement with prior studies (Grant et al., 2019; Hartmann et al., 2015; Hill et al., 2015), distinct variations in mural cell morphology were observed in different vascular segments. Endothelial cells of the parenchymal arteriole are encased by a continuous circumferential layer of vascular smooth muscle cells (Figure 3B), whereas post-arteriole transitional segments (i.e., segments between parenchymal arterioles and the dense capillary region) are covered by “ensheathing pericytes” that are morphologically distinct from vascular smooth muscle cells (Grant et al., 2019; Hartmann et al., 2015; Hill et al., 2015) (Figure 3C). Mural cell coverage of endothelial cells of the capillary bed was incomplete, such that only a few capillary pericytes or thin-stranded capillary processes (Grant et al., 2019; Hartmann et al., 2015; Hill et al., 2015) were observed in our isolated *ex vivo* preparations (Figure 3D). We next compared the propagation velocity of TRPA1-induced conducted vasodilator signals in vascular segments completely covered by mural cells (arterioles and post-arteriole transitional segments) with that in sparsely covered capillaries. The propagation velocity of the vasodilator signal is defined here as the time interval between stimulation of capillaries and start of dilation of the upstream arteriole, normalized to the distance between the site of stimulation and the point where arteriole dilation was measured. We found that the propagation velocity of signals initiated by stimulation of TRPA1 channels in capillaries was significantly slower than that elicited by stimulation of K_ir_ channels (Figure 3E). These data imply that TRPA1 channels in brain capillary endothelial cells mediate a signal that dilates upstream arterioles through a mechanism distinct from that initiated by K_ir_ channels. An analysis of TRPA1-dependent responses in different segments of the cerebral microvasculature showed that the propagation velocity of vasodilator signals from the post-arteriole transitional region (first vascular segment with mural cell coverage) to the upstream parenchymal arteriole was comparable to that of signals produced in this segment by picospritzing capillaries with elevated [K^+^] (Figure 3F). Consequently, we infer that the signal produced by activation of TRPA1 channels propagates through the capillary network from the site of initiation to the transitional region more slowly than the fast-electrical signal initiated by activation of K_ir_ channels. We propose a biphasic propagation model of conducted vasodilator signaling in which activation of TRPA1 channels in capillary endothelial cells generates a slowly propagating intercellular signal that is converted to a rapidly conducted electrical signal in vascular segments fully covered by mural cells (Figure 3G).

**Figure 3.**
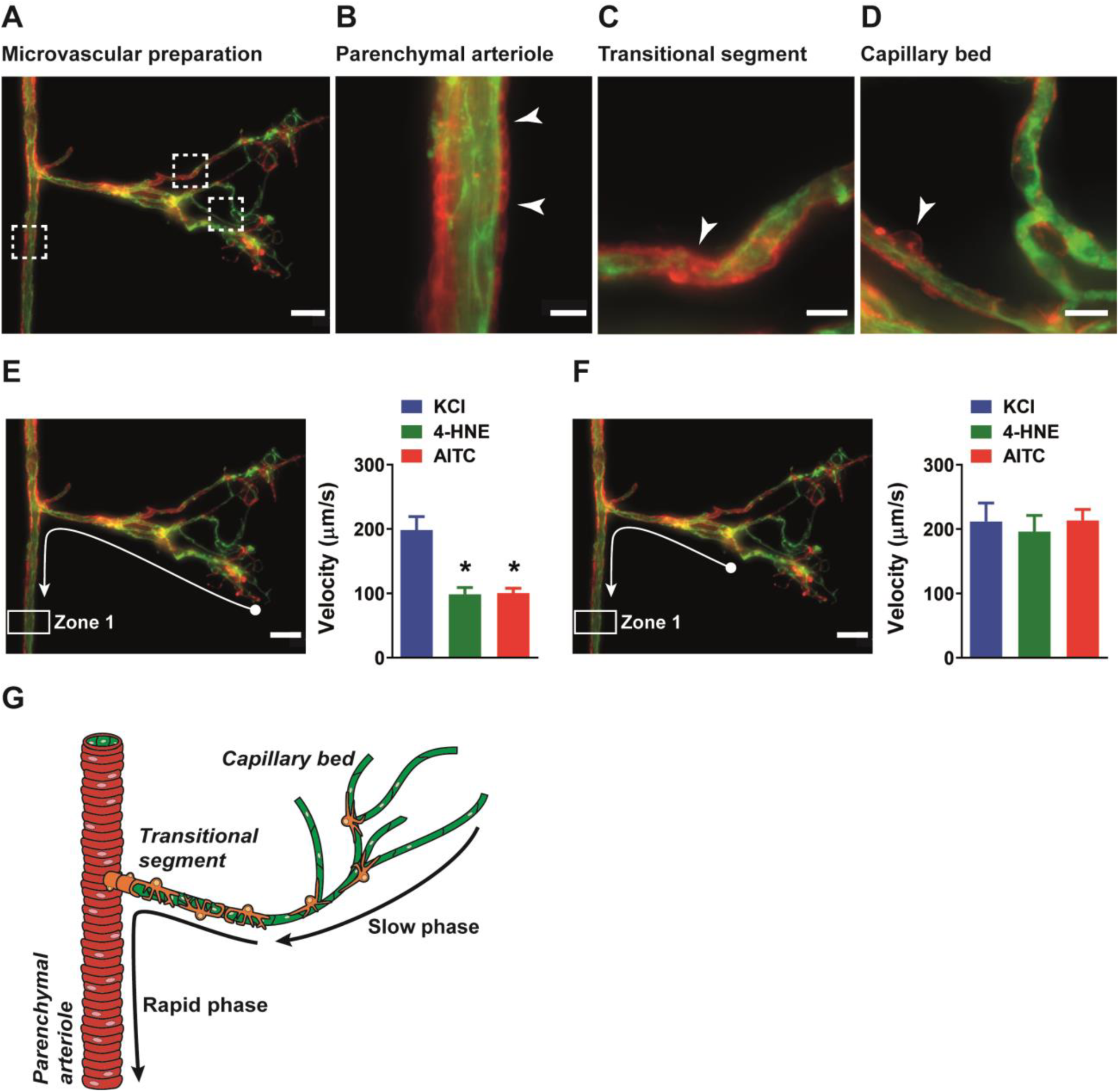
Biphasic velocity of conductive vasodilation following activation of capillary TRPA1 channels. A) Representative fluorescence image of an *ex vivo* microvascular preparation obtained from *VEC-cre mT/mG* mice. Endothelial cells express membrane bound EGFP (green) and all other cell types express tdTomato (red). Scale bar = 50 µm. B to D) Magnified images of the parenchymal arteriole (B), post-arteriole transitional segment (C) and capillaries (D) showing differential mural cell coverage (white arrow) in these vascular segments. Scale bars = 10 µm. E) *Left*: Conduction velocity of the vasodilator signal was calculated from the time interval and distance traveled to a specific point in the upstream arteriole (Zone 1, white arrow) following application of drugs onto capillary extremities (white ball). *Right*: Responses to the TRPA1 channel activators AITC (30 µM) and 4-HNE (1 µM) were significantly slower than the response to activation of K_ir_ channels with KCl (10 mM) (n = 6 to 15 preparations from 4 to 7 animals; *P < 0.05). F). *Left:* Velocity measurements were calculated from the post-arteriole transitional segment (adjacent to the capillary bed, white ball) to a point in the upstream arteriole (Zone 1, white arrow). *Right:* Velocity following AITC (30 µM) or 4-HNE (1 µM) treatment was comparable to that following application of KCl (60 mM) (n = 6 preparations from 4 animals). G) Proposed model showing that activation of TRPA1 channels on capillaries produces a signal that slowly propagates through the capillary network and a faster propagating signal in arterioles and the post-arteriole transitional region.

### Slow-phase propagation of vasodilator signals through the capillary network requires Ca^2+^ signals generated by endothelial cell Panx1 channels and ATP

Our next goal was to identify the mechanisms underlying slow intercellular propagation of vasodilator signals initiated by TRPA1 channels within regions of the capillary network more distal to the feeding arteriole. Pannexin proteins are centrally important for intercellular signaling (Barbe, Monyer, & Bruzzone, 2006). Accordingly, we probed the role of pannexins in conducted vasodilator responses using cerebral microvascular preparations from tamoxifen-inducible, endothelial cell-specific *Panx1*-knockout (*Panx1*-ecKO) mice (Lohman et al., 2015). These mice were generated by crossing mice homozygous for *loxP* sites flanking exon 3 of *Panx1* (*Panx1^fl/fl^*) with mice heterozygous for inducible vascular endothelial cadherin (*VECad*)*-cre*. We found that dilation of upstream parenchymal arterioles following focal application of 4-HNE or AITC to capillary extremities was significantly blunted in *ex vivo* preparations obtained from *Panx1*-ecKO mice compared with those from tamoxifen-injected control mice expressing only *VECad-cre* as well as vehicle (peanut oil)-injected *Panx1*-ecKO mice (Figure 4A to C). In contrast, conducted vasodilation initiated by stimulation of K_ir_ channels in brain capillary endothelial cells was not affected by endothelial cell *Panx1* knockout (Figure 4D). These data demonstrate that TRPA1 channels in brain capillary endothelial cells initiate conducted vasodilator responses through an endothelial cell Panx1-dependent mechanism that is independent of the response initiated by K_ir_ channels.

**Figure 4.**
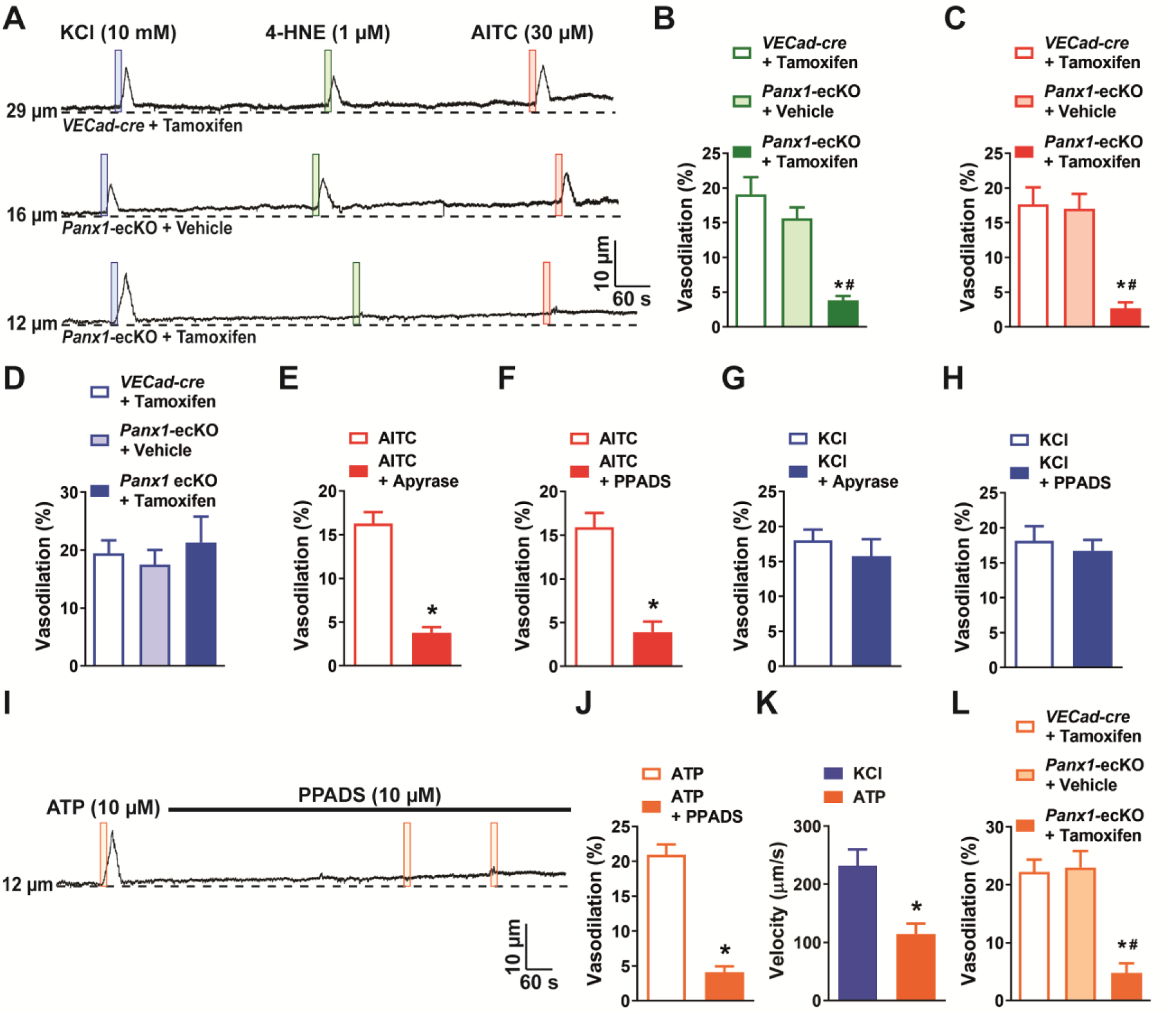
Activation of capillary TRPA1 channels produces a purinergic signal that travels through the capillary bed. A) Representative traces showing that application of AITC (30 µM; red box) or 4-HNE (1 µM; green box) onto capillary extremities did not dilate the upstream arteriole in microvascular preparations from *Panx1*-ecKO mice, whereas vasodilatory effects of elevated KCl (10 mM; blue box) were unchanged. B and C) Summary data showing that neither 4-HNE (1 µM) (B) nor AITC (30 µM) (C) evoked dilation of upstream arterioles in preparations from *Panx1*-ecKO mice (n = 6 preparations from 5 animals; *P < 0.05 vs. *VECad-cre* injected with tamoxifen; ^#^P < 0.05 vs. *Panx1*-ecKO mice injected with vehicle (peanut oil)). D) Summary data showing that the response to elevated KCl (10 nM) was unchanged in preparations from *Panx1*-ecKO mice (n = 6 preparations from 5 animals). E) Summary data showing that catabolism of extracellular purines with apyrase (1 U/mL) blunted the response to AITC (30 µM) in microvascular preparations from wild-type mice (n = 9 preparations from 8 animals; *P < 0.05). F) Summary data showing that the pan-P_2_X inhibitor PPADS (10 µM) inhibited the response to AITC (30 µM) (n = 9 preparations from 6 animals; *P < 0.05). G and H). The response to elevated KCl (10 mM) was unaffected by treatment with apyrase (1 U/mL) (G) or PPADS (10 µM) (H) (n = 9 preparations from 6 to 8 animals). I) Representative trace showing that application of ATP (10 µM; orange box) onto capillary extremities dilated the upstream arteriole in microvascular preparations from wild-type mice, and that this response was blocked by the pan-P_2_X inhibitor PPADS (10 µM). J) Summary data showing that PPADS (10 µM) inhibited the response ATP (10 µM) responses (n = 6 preparations from 4 animals; *P < 0.05). K) The velocity of the ATP (10 µM) response was significantly slower compared with that to activation of K_ir_ channels with KCl (60 mM) (n = 6 preparations from 4 animals; *P < 0.05). L) Summary data showing that ATP (10 µM) did not evoke dilation of upstream arterioles in preparations from *Panx1*-ecKO mice (n = 6 preparations from 3 to 4 animals; *P < 0.05 vs. *VECad-cre* injected with tamoxifen; ^#^P < 0.05 vs. *Panx1*-ecKO mice injected with vehicle (peanut oil)).

Prior studies have demonstrated that increases in intracellular [Ca^2+^] cause the release of ATP through Panx1 channels (Dahl, 2015). Released ATP subsequently activates Ca^2+^-permeable ionotropic purinergic (P_2_X) receptors on adjacent cells to increase intracellular Ca^2+^, providing a means for the intercellular propagation of Ca^2+^ signals (Barbe et al., 2006; Locovei, Wang, & Dahl, 2006; Suadicani, Iglesias, Spray, & Scemes, 2009; Vanlandewijck et al., 2018; Zsembery et al., 2003). We used a pharmacological approach to determine if this mechanism was responsible for the propagation of conducted vasodilator signals in the brain capillary network. Pre-incubation of microvascular preparations with apyrase (1 U/mL) to catabolize extracellular purines (Figure 4E) and administration of the pan-P_2_X inhibitor PPADS (10 µM) blocked the dilation of upstream parenchymal arteriole segments in response to application of AITC to capillaries (Figure 4F). However, apyrase and PPADS had no effect on conducted vasodilation initiated by stimulation of brain capillary endothelial cell K_ir_ channels with 10 mM K^+^ (Figure 4G and H). We also found that direct application of ATP onto capillary beds stimulated conducted dilation in preparations from wild-type mice, an effect that was blocked by PPADS (Figure 4I and J). The velocity of ATP-conducted responses was slower than that of responses initiated by stimulation of K_ir_ channels and was comparable to that of signals produced by stimulation of TRPA1 channels (Figure 4K). Consistent with these findings, application of ATP onto capillaries did not initiate conducted vasodilation in *ex vivo* preparations from *Panx1*-ecKO mice (Figure 4L). These data support the concept that stimulation of TRPA1 channels on capillary endothelial cells initiates the release of ATP through endothelial cell Panx1 channels. Released ATP, in turn, stimulates P_2_X receptors on adjacent cells to generate a propagating Ca^2+^ signal.

We then asked, how far does the intercellular signal initiated by TRPA1 channels travel within the capillary network? To this end, we determined the length of cerebral capillary endothelial cells by measuring the distance between nuclei in isolated capillary networks labeled with isolectin B4 and DAPI. We found that the average capillary endothelial cell length was 28.7 ± 0.6 µm (Supplemental Figure 4A and B). Given that the length of the capillary segment in our *ex vivo* microvascular preparation is in the range of 100 to 150 µm (Supplemental Figure 4C), we estimate that the signal initiated by TRPA1 channels propagates through 3 to 5 endothelial cells in the capillary network before reaching post-arteriole transitional segments with mural cell coverage.

TRPA1 channels conduct mixed cation currents with a high Ca^2+^ fraction (Nilius, Prenen, & Owsianik, 2011), and we have previously detected distinct TRPA1 channel-mediated Ca^2+^-signaling events in cerebral endothelial cells (Sullivan et al., 2015). Ca^2+^-imaging studies performed using microvascular preparations obtained from transgenic mice expressing the genetically encoded Ca^2+^ indicator GCaMP8, exclusively in the endothelium (*VEC-GCaMP8*) (Ohkura et al., 2012) revealed that focal application of AITC onto the distal extremities of capillaries initiated an increase in intracellular [Ca^2+^] (Supplemental Movie 3). Moreover, this increase in intracellular [Ca^2+^] triggered similar increases in neighboring regions, indicating that the Ca^2+^ signal propagates through the capillary network (Figure 5A). AITC-induced Ca^2+^ signals were abolished by superfusing the preparation with HC-030031 (Supplemental Movie 4, Figure 5B and C), suggesting that initiation of the signal required the activity of TRPA1 channels. As our model predicts that the propagation velocity of the vasodilatory signal within the capillary network is slow, we therefore determined the velocity at which Ca^2+^ signals travels through the capillary network. The propagation velocity of the Ca^2+^ signal is defined here as the time interval between the start of an increase in intracellular [Ca^2+^] at one region of interest and the start of an increase in intracellular [Ca^2+^] in a neighboring region, normalized to the distance between the two regions. As expected, the velocity of vasodilator Ca^2+^ signals within the capillaries travel at speeds slower than the rapid vasodilator signal travelling through vascular segments covered by mural cells (Figure 5D). Focal application of ATP was also found to initiate an increase in intracellular [Ca^2+^], that propagates to adjacent cells (Supplemental Movie 5, Figure 5E). PPADS abolished ATP-induced Ca^2+^ signal, suggesting that initiation and propagation of Ca^2+^ signals require P_2_X receptors (Supplemental Movie 6, Figure 5F and G). The velocity of ATP-induced Ca^2+^ signals is similar to that produced by activating TRPA1 channels, suggesting a convergent vasodilator signaling pathway (Figure 5H). The duration, rise time and decay time of the Ca^2+^ signal evoked by AITC and ATP were also comparable (Supplemental Figure 5 A to C). We next validated that the vasodilator signal through the capillaries requires influx of extracellular Ca^2+^ through purinergic P_2_X receptors by superfusing preparations with aCSF devoid of extracellular Ca^2+^. We found that ATP-induced Ca^2+^ signals were abolished in preparations under 0 Ca^2+^ conditions, however the response returned once extracellular Ca^2+^ (2 mM) was reintroduced (Supplemental Movie 7 to 9, Figure 5I and J). This suggests that the propagative Ca^2+^ signal is dependent on ionotropic purinergic P_2_X receptor, and not through the release of Ca^2+^ from intracellular stores downstream of G protein-coupled P_2_Y receptor activation.

**Figure 5.**
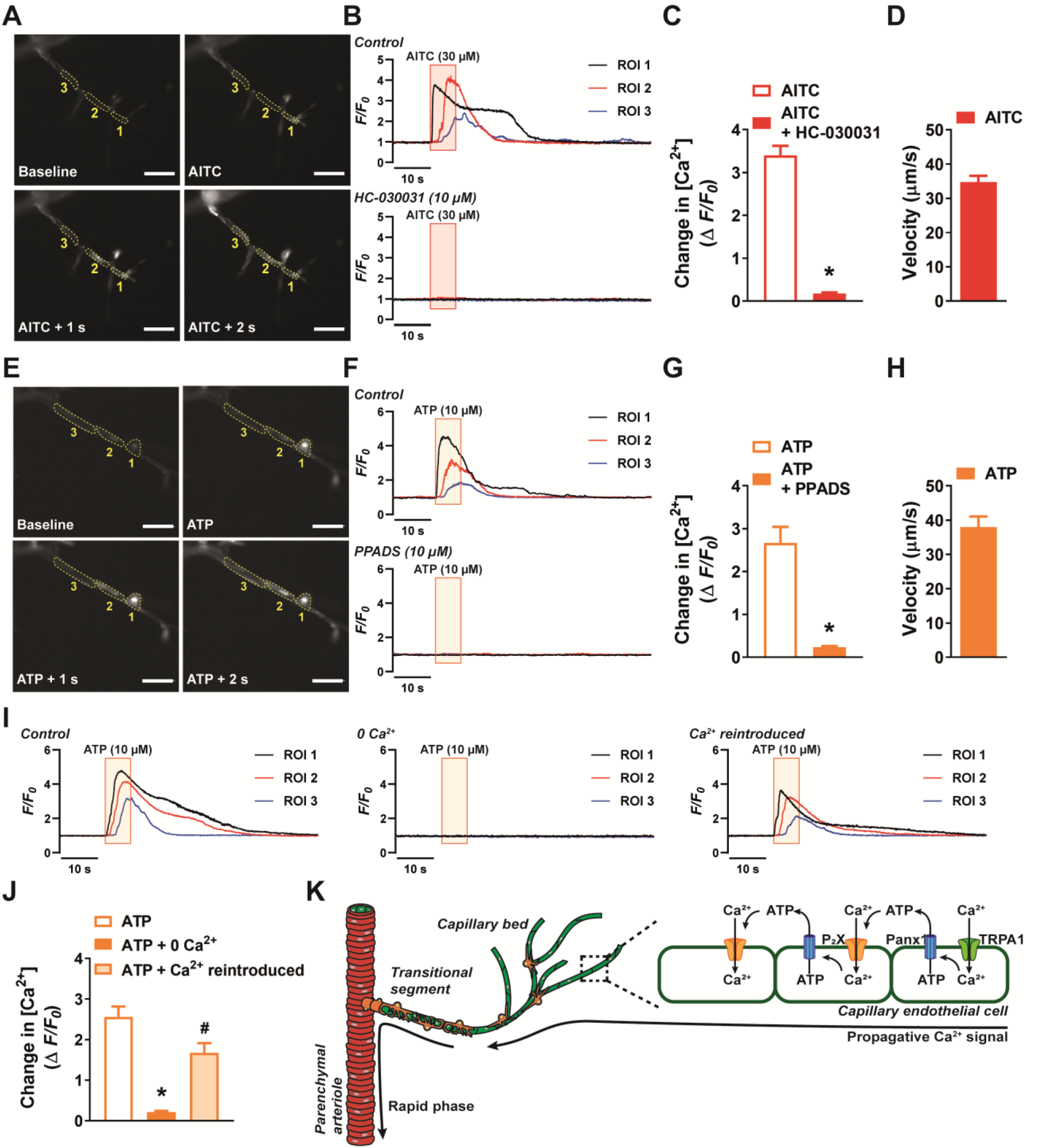
Activation of capillary TRPA1 channels and purinergic receptors produces a Ca^2+^ response that travels through the capillary bed. A) Representative time course images demonstrating the fractional increase in fluorescence (F/F_0_) of the Ca^2+^ signal in a capillary segment in microvascular preparations from transgenic mice expressing a genetically encoded Ca^2+^ indicator, GCaMP8, exclusively in the endothelium (*VEC-GCaMP8*). Focal application of AITC (30 µM) to distal capillaries evoked a propagative Ca^2+^ signal that was observed in adjacent region of interests. Scale bar = 50 µm. B) Representative traces showing the fractional increase in florescence following application of AITC (30 µM; red box) that was blocked by the selective TRPA1 antagonist HC-030031 (10 µM). C) Summary data showing that HC-030031 (10 µM) inhibited the response to AITC (30 µM) (n = 6 preparations from 5 animals; *P < 0.05). D) The velocity of the response through capillary segments evoked by activation of TRPA1 channels (n = 6 preparations from 5 animals). E) Representative time course images demonstrating the fractional increase in fluorescence of the Ca^2+^ signal produced following focal application of ATP (10 µM) to distal capillaries. Propagation of the Ca^2+^ signal was observed in the adjacent region of interests following ATP (10 µM) application. Scale bar = 50 µm. F) Representative traces showing the fractional increase in florescence following application of ATP (10 µM; orange box) that was blocked by the pan-P_2_X inhibitor PPADS (10 µM). G) Summary data showing that PPADS (10 µM) inhibited the response to ATP (10 µM) (n = 7 preparations from 7 animals; *P < 0.05). H) The velocity of the response through capillary segments evoked by activation of purinergic receptors (n = 13 preparations from 12 animals). I) Representative traces showing the fractional increase in florescence following application of ATP (10 µM; orange box) was abolished when the preparation was superfused in Ca^2+^-free aCSF. The response was restored following the reintroduction of extracellular Ca^2+^. J) Summary data showing that bathing the preparation in Ca^2+^-free aCSF solution attenuated the response to ATP (10 µM), but was restored once Ca^2+^ (2 mM) was reintroduced (n = 6 preparations from 5 animals; *P < 0.05 vs. control response; ^#^P < 0.05 vs. preparations bathed in 0 Ca^2+^). K) Illustration of the proposed model for signal propagation through the capillary bed. Increases in intracellular [Ca^2+^] caused by TRPA1 channel-mediated Ca^2+^ influx induce ATP release through Panx1 channels, which in turn activates purinergic P_2_X receptors on the adjacent endothelial cell.

Taken together, these data support a model in which slow, short-range intercellular Ca^2+^ signals propagate through capillary endothelial cells via a mechanism that is dependent on ATP release through Panx1 channels and activation of purinergic P_2_X receptors on adjacent endothelial cells (Figure 5K). This pathway is necessary for the dilation of upstream arterioles in response to stimulation of TRPA1 channels in capillary endothelial cells.

### Rapid-phase propagation of vasodilator signals is initiated in the post-arteriole transitional region and requires K_ir_ channel activity

We next turned to the mechanisms responsible for the rapid phase of the propagating vasodilator signal. Because signal propagation velocities were the same in post-arteriole transitional segments and parenchymal arterioles, regardless of the initiating stimulus, we hypothesized that propagation in this segment occurs by rapid electrical communication, as previously described by Longden et al. (Longden et al., 2017). To test this, we performed experiments using a modified capillary-arteriole microvascular preparation in which the capillary tree was removed while leaving the post-arteriole transitional segment intact (Figure 6A). In control experiments, we found that Evans Blue dye ejected from a picospritzer near the post-arteriole transitional segment did not spread to the upstream parenchymal arteriole (Supplemental Movie 10), supporting our ability to selectively stimulate the transitional segments.

**Figure 6.**
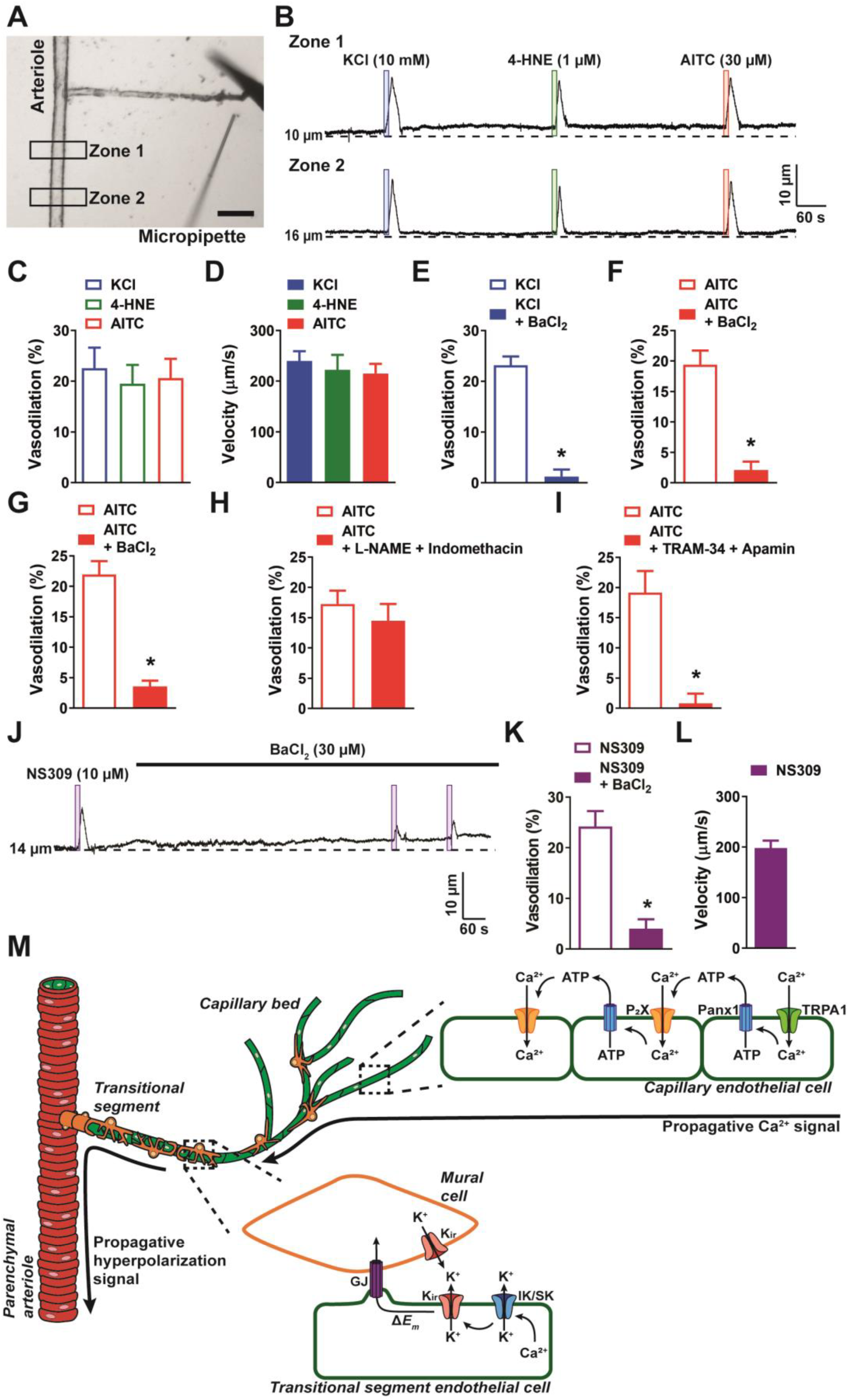
Rapid propagation of TRPA1-induced vasodilation is dependent on K_ir_, IK, and SK channels. A) Representative image of a cerebral microvascular preparation in which the capillary bed was removed while leaving the post-arteriole transitional segment intact. Scale bar = 100 µm. A picospritzing cannula was used to directly apply drugs to the post-arteriole transitional segment. B) Representative traces showing that application of elevated KCl (10 mM; blue box), AITC (30 µM; red box), or 4-HNE (1 µM; green box) onto the post-arteriole transitional segment increased the lumen diameter of the upstream arteriole in microvascular preparations from wild-type mice. C) Summary data showing dilation produced by elevated KCl (10 mM), AITC (30 µM) and 4-HNE (1 µM) (n = 6 to 8 preparations from 4 to 5 animals). D) Velocity was calculated from the post-arteriole transitional segment to a point in the upstream arteriole following localized drug application onto the post-arteriole transitional segment. Velocity measurements were comparable for elevated KCl (10 mM), 4-HNE (1 µM) and AITC (30 µM) (n = 6 to 8 preparations from 4 to 5 animals). E and F) Summary data showing that BaCl_2_ (30 µM) significantly reduced the response to elevated KCl (10 mM) (E) and AITC (30 µM) (F) when applied to the post-arteriole transitional segment (n = 6 preparations from 4 animals; *P < 0.05). G) Summary data showing that BaCl_2_ (30 µM) significantly attenuated the response to AITC (30 µM) when directly applied onto capillary extremities (n = 6 preparations from 4 animals; *P < 0.05). H) Summary data showing that combined inhibition of nitric oxide synthase and cyclooxygenase with L-NAME (100 µM) and indomethacin (10 µM), respectively, did not affect the response to AITC (30 µM) when directly onto capillary extremities (n = 7 preparations from 4 animals; *P < 0.05). I) Summary data showing that combined IK and SK channel inhibition with TRAM-34 (1 µM) and apamin (300 nM), respectively, significantly attenuated the response to AITC (30 µM) when directly applied onto capillary extremities (n = 7 preparations from 5 animals; *P < 0.05). J and K) Representative trace (J) and summary data (K) showing that application of the IK and SK channel activator NS309 (10 µM; purple box) directly onto the post-arteriole transitional segment dilated the upstream arteriole, and that this response was attenuated by inhibition of K_ir_ channels with BaCl_2_ (30 µM) (n = 6 preparations from 4 animals; *P < 0.05). L) The velocity of the response produced by application of the IK and SK channel activator NS309 (10 µM) onto the post-arteriole transitional segment was similar to that produced by activation of K_ir_ channels or TRPA1 channels. M) Illustration of the proposed signaling model depicting events following activation of capillary endothelial TRPA1 channels. Once the Ca^2+^ signal from the capillary bed arrives at the post-arteriole transitional segment, IK and SK channels are activated and facilitate K^+^ efflux. This in turn activates K_ir_ channels, which propagate the signal retrogradely to cause dilation of the upstream arteriole.

Interestingly, we found that focal application of elevated [K^+^], 4-HNE, or AITC onto the post-arteriole transitional segment caused dilation of the upstream parenchymal arteriole (Figure 6B and C), indicating that functional K_ir_ and TRPA1 channels are present in these transitional segments and are capable of producing conducted vasodilator responses when stimulated. The propagation velocities of the conducted vasodilator signals produced by stimulating post-arteriole transitional segment with elevated [K^+^], AITC, or 4-HNE were nearly identical, suggesting a common signal-propagation mechanism (Figure 6D). We also found that vasodilator responses to elevated [K^+^] (Figure 6E) and AITC (Figure 6F) were blocked by inhibiting K_ir_ channels with BaCl_2_. The rapid signal-propagation velocity and sensitivity to blockade by BaCl_2_ suggest that fast electrical communication involving K_ir_ channels (Longden et al., 2017) is a convergent means of vasodilator signal conduction in vascular segments with mural cell coverage. We also found that superfusing the *ex vivo* preparation with BaCl_2_ significantly blunted the response to localized application of AITC onto capillary extremities (Figure 6G). Interestingly, focal application of ATP onto the post-arteriole transitional segment did not evoke dilation of upstream arterioles (Supplemental Figure 6A and B), suggesting that ATP-dependent propagation must be initiated at more distal points in the capillary network.

We next investigated how the slowly propagating Ca^2+^ signals initiated by capillary endothelial cell TRPA1 channels are converted to rapidly propagating electrical signals in mural cell-covered post-arteriole transitional segments to cause dilation of upstream parenchymal arterioles. Combined inhibition of nitric oxide synthase with L-NAME (100 µM) and cyclooxygenase with indomethacin (10 µM) in microvascular preparations had no effect on arteriole dilation induced by stimulation of capillary extremities with AITC (Figure 6H). However, inhibition of IK and SK channel activity by superfusing TRAM34 (1 µM) and apamin (300 nM), respectively, significantly blunted upstream dilation in response to focal stimulation of capillary endothelial cell TRPA1 channels (Figure 6I), demonstrating that IK and SK channel activity is necessary for the conducted vasodilator responses initiated by brain capillary endothelial cell TRPA1 channels. Although it has been reported that functional IK and SK channels are not present in capillary endothelial cells (Longden et al., 2017) (Supplemental Figure 1), our functional data suggest that IK and SK channels are expressed in post-arteriole transitional segments, where they convert slowly propagating intercellular Ca^2+^ signals initiated by capillary TRPA1 channels into rapidly propagating electrical signals. Direct application of NS309 (10 µM) directly onto parenchymal arterioles, which are known to express endothelial IK and SK channels caused dilation, whereas focal stimulation of capillary extremities did not (Supplemental Figure 7A and B). Surprisingly, we found that focal application of NS309 directly onto the post-arteriole transitional segment induced dilation of the upstream parenchymal arteriole, indicating that IK and SK channels are expressed in this region and that their activation results in conducted vasodilation (Figure 6J and K). We also found that this response was inhibited by BaCl_2_ (Figure 6K) and propagated at the same velocity as signals initiated by stimulating the post-arteriole transitional segment with elevated [K^+^] and TRPA1 agonists (Figure 6L). These data support the concept that the Ca^2+^ signals arriving from the capillary network activate IK and SK channels within the post-arteriole transitional segment to initiate a fast-conducting vasodilator response that requires K_ir_ channel activity (Figure 6M).

### Stimulation of brain endothelial cell TRPA1 channels increases local blood flow ***in vivo*.**

We next performed experiments to determine if stimulation of endothelial cell TRPA1 channels increase localized cerebral blood flow *in vivo*. Red blood cell (RBC) flux within capillaries and changes in diameter of upstream arterioles were visualized through a cranial window using two-photon laser-scanning microscopy (Figure 7A). Fluorescein isothiocyanate (FITC)-conjugated dextran (*i.v.*) was administered to animals to allow visualization of the vasculature and contrast imaging of RBCs. TRPA1 channels were locally activated by pressure ejecting AITC directly onto the capillary bed using a micropipette. In control *Trpa1^fl/fl^* animals, local application of AITC (30 µM) onto capillaries produced a significant increase in RBC flux within the stimulated capillary bed (Figure 7B and C). The latency of this response (20.6 ± 4.0 s, n = 11) was slow compared with the response induced by applying elevated [K^+^] (3.8 ± 0.9 s), as reported by Longden et al. (Longden et al., 2017). This finding is consistent with the slower kinetics of the TRPA1-mediated vasodilator response compared with the K_ir_-mediated response observed in our *ex vivo* microvascular preparations (Figure 3A). In contrast, AITC failed to elicit an increase in RBC flux in almost all (6 of 7) *Trpa1*-ecKO mice tested (Figure 7D to F). We also confirmed vasodilatory responses to capillary stimulation by assessing changes in the diameter of feeding arterioles, demonstrating that local application of AITC onto capillaries significantly increased the diameter of upstream arterioles in *Trpa1^fl/fl^* animals (Figure 7G). Furthermore, the increase in diameter observed in arterioles of *Trpa1^fl/fl^* animals was greater than that observed in *Trpa1*-ecKO mice (*Trpa1^fl/fl^* mice 2.39 ± 0.34 µm (n = 11) vs. *Trpa1*-ecKO mice 0.47 ± 0.16 µm (n = 7), P < 0.05) (Figure 7H and I). A small increase in diameter was detected in *Trpa1*-ecKO mice, possibly due to involvement of TRPA1 channels in other cell types, such as astrocytes (Shigetomi, Tong, Kwan, Corey, & Khakh, 2011). These data indicate that activation of endothelial cell TRPA1 channels in the brain dilates upstream arterioles and increases RBC flux in capillaries.

**Figure 7.**
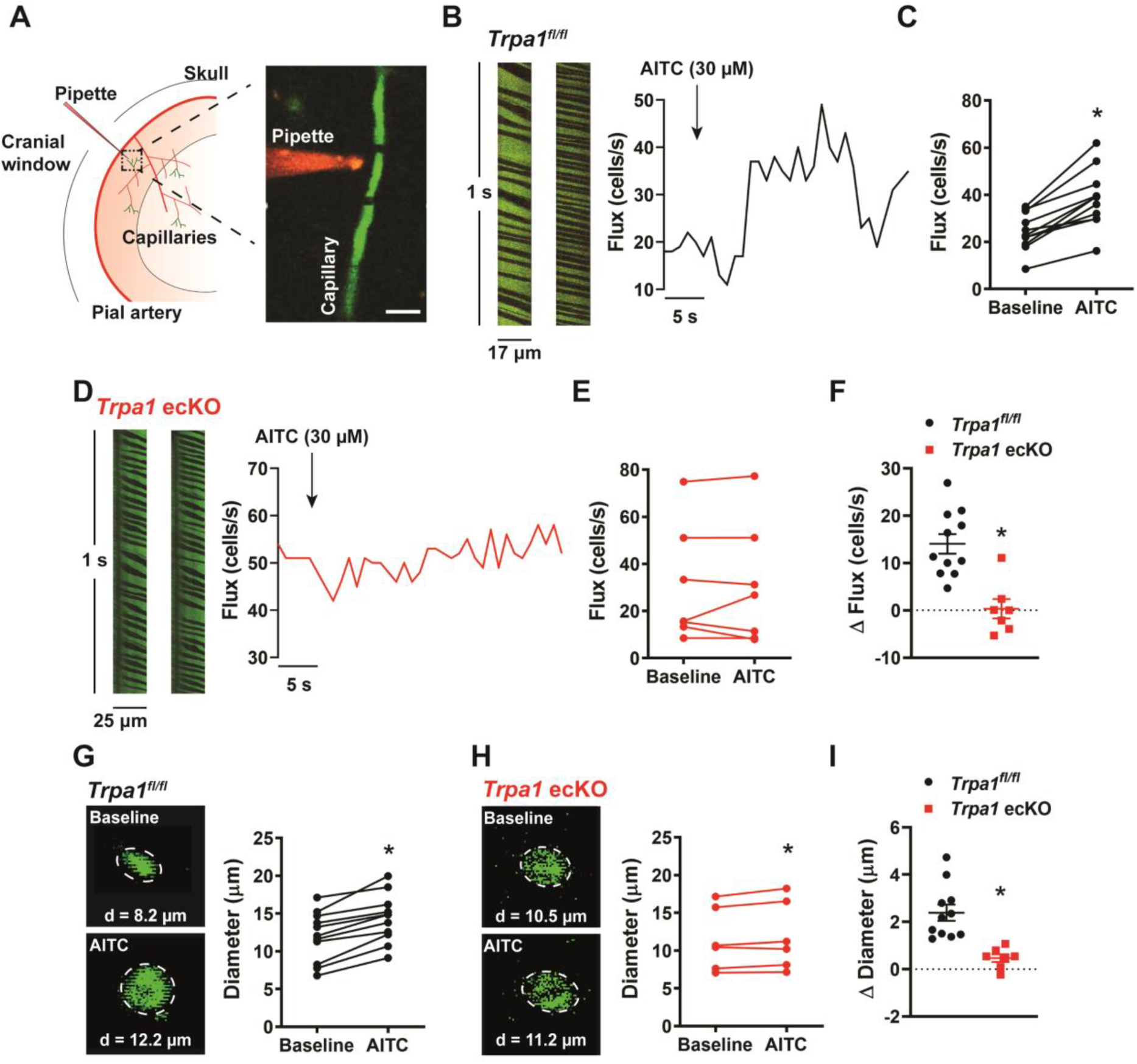
In vivo stimulation of capillary TRPA1 channels increases RBC flux within the capillary bed. A) Experimental illustration. Mice were injected with FITC-conjugated dextran to allow visualization of the vasculature through a cranial window using two-photon laser-scanning microscopy. A pipette containing AITC and TRITC-dextran was positioned next to a capillary. Scale bar = 10 µm. B) RBC flux through the capillary bed was examined *in vivo* using two-photon imaging. TRPA1 channels on capillary endothelial cells were locally stimulated by picospritzing AITC (30 µM) directly onto a capillary bed. Representative line-scan plots of a capillary (*left*) and representative time-flux trace (*right*) demonstrating changes in RBC flux before and after application of AITC (30 µM) in control *Trpa1^fl/fl^* mice. C) Summary data showing that AITC (30 µM) increased RBC flux in *Trpa1^fl/fl^* mice (n = 11 animals; *P < 0.05). D) Representative line-scan plots of a capillary (*left*) and representative time-flux trace (*right*) demonstrating that application of AITC (30 µM) had no effect in *Trpa1*-ecKO mice. E) Summary data showing that AITC (30 µM) had no effect in *Trpa1*-ecKO mice (n = 7 animals). F) Change in RBC flux in control *Trpa1^fl/fl^* versus *Trpa1*-ecKO mice (n = 7 to 11 animals; *P < 0.05). G and H) Representative images and summary data showing change in diameter of upstream arterioles following application of AITC (30 µM) onto the capillary bed of control *Trpa1^fl/fl^* (G) and *Trpa1*-ecKO (H) mice (n = 7 to 11 animals; *P < 0.05). I) Change in arteriole diameter was significantly greater in *Trpa1*-ecKO mice compared to control *Trpa1^fl/fl^* (n = 7 to 11 animals; *P < 0.05).

### Functional hyperemia in the somatosensory cortex requires endothelial cell TRPA1 channels

To assess the role of TRPA1 channels in functional hyperemia *in vivo*, we measured relative changes in blood flow in the somatosensory cortex using a thinned-skull mouse model (Figure 8A). Changes in blood flow were induced by contralateral whisker stimulation and were recorded using laser-Doppler flowmetry (Girouard et al., 2010; Park et al., 2014). Whisker stimulation reliably and reproducibly increased cerebral blood flow in *Trpa1^fl/fl^* mice (Figure 8B). Control experiments indicated that this was not due to artifactual noise, as stimulation of ipsilateral whiskers was without effect (Supplemental Figure 8A and B). To determine if TRPA1 channels are involved in this functional hyperemic response, we treated mice with HC-030031 (100 mg/kg, *i.p.* for 30 min). HC-030031 treatment significantly attenuated the increase in cerebral blood flow following whisker stimulation (Figure 8B). To demonstrate the contribution of endothelial cell TRPA1 channels, we assessed functional hyperemia in *Trpa1*-ecKO mice. The basal increase in blood flow following whisker stimulation was significantly blunted (∼40% less) in *Trpa1*-ecKO mice compared with *Trpa1^fl/fl^* mice (Figure 8C). Collectively, these data demonstrate that endothelial cell TRPA1 channels are necessary for functional hyperemia in the somatosensory cortex of the brain.

**Figure 8.**
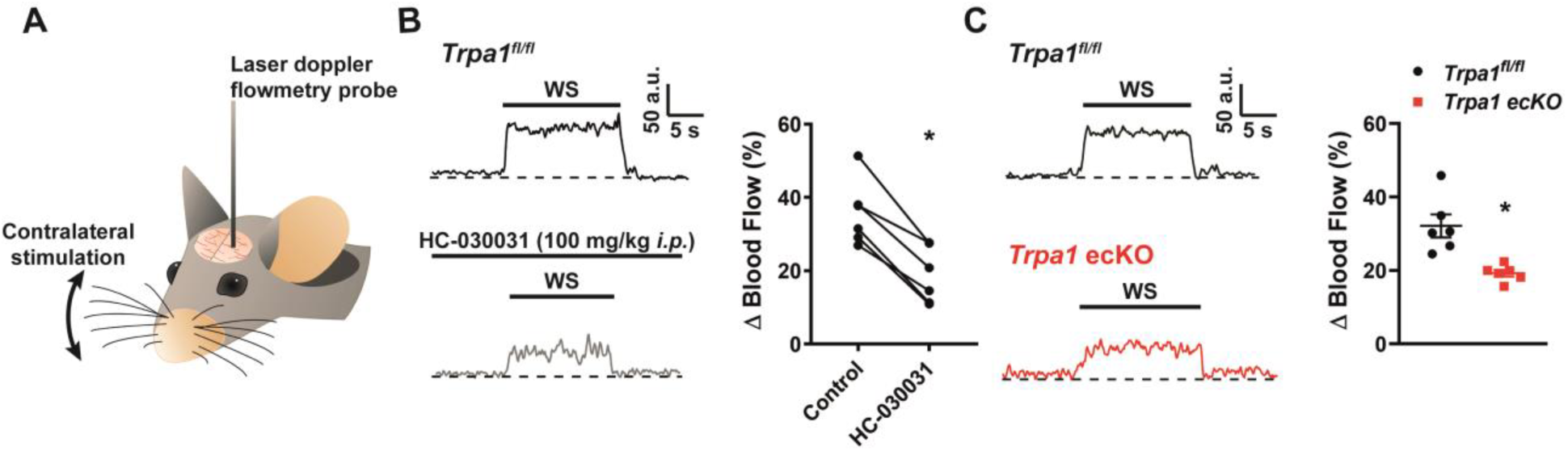
Functional hyperemia is dependent on brain capillary TRPA1 channels. A) Experimental illustration. Functional hyperemia was assessed in the somatosensory cortex through a thinned skull. Relative changes in blood flow in response to contralateral whisker stimulation were recorded using laser-Doppler flowmetry. B) Representative traces (*left*) and summary data (*right*) showing the hyperemic response in the somatosensory cortex following contralateral whisker stimulation (WS), measured using laser-Doppler flowmetry in control *Trpa1^fl/fl^* mice. Treatment with HC-030031 (100 mg/kg *i.p.*) reduced the hyperemic response (n = 6 animals). C) Representative traces (*left*) and summary data (*right*) showing the hyperemic response in the somatosensory cortex following contralateral whisker stimulation (WS) was reduced in *Trpa1*-ecKO mice (n = 6 animals).

## Discussion

The parenchyma of the brain is exposed to an ever-changing milieu of physical, environmental, endocrine, paracrine, metabolic, and neurochemical stimuli. The neurovascular apparatus must detect and decode signals generated by active areas to properly direct regional blood flow and ensure optimal brain function. The preponderance of the brain vasculature is composed of densely packed capillary networks; an optical volumetric analysis has revealed that the soma of all central neurons are located no more than 15 μm from a brain capillary endothelial cell (Tsai & Kleinfeld, 2009). Building on these observations, Longden et al. introduced a convincing new paradigm for NVC, demonstrating that conducted vasodilator responses orchestrated by capillary endothelial cells provoke functional hyperemia in the brain through propagating electrical (hyperpolarizing) signals initiated and sustained by K_ir_ channels. Thus, the brain’s extensive capillary network serves as a sensory web for detecting [K^+^] released at active neuronal synapses (Longden et al., 2017). The NVC response is essential for life itself, and the multiplicity of distinct signaling modalities in the active brain imply that brain capillary endothelial cells possess an equally broad assortment of complementary and overlapping sensors. Here, we investigated this concept, providing evidence that activation of brain capillary endothelial cell TRPA1 channels promotes dilation of upstream parenchymal arterioles through a novel, biphasic intercellular signaling mechanism that is dependent on Panx1 channels. Our findings also identify purinergic receptors on capillary endothelial cells as part of this signal conduction pathway and as independent detectors of released ATP. We further demonstrate that focal stimulation of K_ir_ and TRPA1 channels in the post-arteriole transitional region results in conducted dilation of upstream arterioles, expanding the brain’s vascular sensory network to include these segments. We conclude that multiple molecular sensors that are critically important for NVC and functional hyperemia in the brain, including TRPA1 channels and purinergic receptors, are present in brain capillary endothelial cells.

Our findings demonstrate that capillary endothelial cell TRPA1 channels initiate signals that propagate through the cerebral microvasculature to cause dilation of upstream parenchymal arterioles, locally increase RBC flux, and produce functional hyperemia in the brain. TRPA1 exhibits a unique pattern of expression within the vasculature: it is present in the endothelium of arteries and arterioles in the brain, but not in other organs, such as the heart, dermis, kidney or mesentery (Earley et al., 2009; Pires & Earley, 2018; Sullivan et al., 2015). Activation of vascular TRPA1 channels produces endothelium-dependent dilation of cerebral arteries and arterioles, a mechanism that is neuroprotective during ischemic stroke (Earley et al., 2009; Pires & Earley, 2018; Sullivan et al., 2015). Lipid peroxidation metabolites generated by reactive oxygen species (ROS), such as 4-HNE, directly activate TRPA1 channels (Andersson, Gentry, Moss, & Bevan, 2008; Taylor-Clark et al., 2008). We have shown that superoxide anions (O_2_^-^), generated by the activity of NADPH oxidase 2 (NOX2) or by mitochondrial respiration under hypoxic and ischemic conditions, endogenously generate 4-HNE, which stimulates TRPA1 channel activity in the cerebral endothelium to cause vasodilation (Pires & Earley, 2018; Sullivan et al., 2015). We propose that extracellular ROS generated by astrocytes could act as endogenous agonists of capillary endothelial cell TRPA1 channels during NVC. One possible source of ROS is astrocytic endfoot processes, which encase brain capillaries and have been shown to express NOX2 and produce ROS (Abramov & Duchen, 2005). Alternatively, extracellular ROS generated by metabolically active neurons proximal to brain capillaries could activate TRPA1 channels. Pericytes and microglia cells are also intimately associated with capillaries in the brain and are potential sources of TRPA1-activating ROS metabolites.

The vasodilator signals initiated by brain capillary endothelial cell TRPA1 channels propagate at about half the speed of those initiated by K_ir_, suggesting that the two signals are conducted by different pathways. Prior work has demonstrated that activation of capillary endothelial cell K_ir_ channels generates a fast-conducting electrical signal that hyperpolarizes smooth muscle cells in upstream arterioles to cause vasodilation (Longden et al., 2017). Our data showed that stimulation of capillary endothelial cell TRPA1 channels produced slowly propagating, short-range intercellular signals that are dependent on endothelial expression of Panx1 channels. Panx1 forms transmembrane channels in cerebral arteriole and capillary endothelial cells (Vanlandewijck et al., 2018). Panx1 channels are normally closed, but when activated by increases in intracellular [Ca^2+^], they release ATP (Dahl, 2015; Locovei et al., 2006). Previous studies using exogenous expression systems have shown that Panx1 channels and Ca^2+^-permeable purinergic P_2_X receptors colocalize on the membrane of *Xenopus* oocytes (Locovei, Scemes, Qiu, Spray, & Dahl, 2007) and urothelial cells (Negoro et al., 2014) to form signaling microdomains that permit intercellular propagation of Ca^2+^ signals. In this mechanism, ATP released via Panx1 channels binds and activates P_2_X receptors on neighboring cells, resulting in Ca^2+^ influx and activation of Panx1 channels, thus establishing a propagating intercellular Ca^2+^ signaling pathway (Barbe et al., 2006; Suadicani et al., 2009). Our data suggest that a similar signaling pathway operates in the cerebral capillary network, providing a novel short-range intercellular communication system between neighboring capillary endothelial cells. Our data further suggest that Ca^2+^ influx through TRPA1 channels is the initiator of this pathway, but it is possible that other Ca^2+^-permeable channels present on capillary endothelial cells can activate this pathway to initiate propagative vasodilation if they form a signaling complex with Panx1.

Purinergic signaling is an important endogenous pathway that regulates many cellular processes, including regulation of vascular tone in multiple vascular beds. Activation of ATP-gated, cation-conducting purinergic P_2_ receptors and G protein-coupled receptors results in dual vasoconstriction and vasodilation responses. Sympathetic nerves surrounding vessels of numerous vascular beds contain ATP, which, upon release, acts primarily on P_2_X_1_ receptors to promote Ca^2+^ influx and subsequent contraction of vascular smooth muscle cells (Burnstock, 2017). Endothelial cells are a significant source of ATP that is released in response to certain stimuli, such as shear stress (Schwiebert, Rice, Kudlow, Taylor, & Schwiebert, 2002). Secreted ATP activates endothelial P_2_X and P_2_Y receptors in an autocrine and/or paracrine manner to produce the endothelium-dependent vasodilators nitric oxide and prostacyclin as well as endothelium-dependent hyperpolarizing factors (Burnstock, 2017). In the cerebral circulation, ATP released from surrounding astrocytes (Kisler, Nelson, Montagne, & Zlokovic, 2017; Pascual et al., 2005) and neurons (Fields & Burnstock, 2006; Kisler et al., 2017) can activate ATP-gated receptors on arteriolar vascular smooth muscle cells, resulting in contraction; it can also induce endothelium-dependent vasodilation through the release of paracrine factors (Chen, Kozberg, Bouchard, Shaik, & Hillman, 2014; Ralevic & Dunn, 2015). Our data support a model in which endogenous ATP derived from astrocytes and/or neurons directly activates P_2_X receptors expressed on capillary endothelial cells to initiate Panx1-dependent conducted vasodilation, providing an additional capillary-based mechanism for initiation of NVC.

Our findings indicate that post-arteriole transitional segments are vital for functional hyperemic responses in the brain. These transitional vascular segments are composed of endothelial cells encompassed by specialized mural cells called ensheathing pericytes (Grant et al., 2019; Hartmann et al., 2015; Hill et al., 2015). Ensheathing pericytes are morphologically distinct from other types of mural cell types as they possess a protruding cell body and are elongated (Grant et al., 2019; Hartmann et al., 2015). Recent reports suggest that ensheathing pericytes are contractile and regulate the diameter of post-arteriole transitional segment to redistribute blood to neuronally active sites (Hill et al., 2015). Our findings suggest that post-arteriole transitional segments serve two additional functions during NVC. First, focal stimulation of these segments with substances that activate K_ir_ or TRPA1 channels initiates conducted dilation of upstream arterioles. These data suggest that the brain’s vascular sensory web extends throughout the capillary tree to include the post-arteriole transitional segment. Interestingly, the mechanism of ATP-induced signal propagation is unique to more distal reaches of the capillary network, as focal application of ATP onto the post-arteriole transitional region failed to dilate upstream arterioles. Secondly, our data show that post-arteriole transitional segments are critically important for the translation of slowly propagating short-range Ca^2+^ signals originating deeper in the capillary bed into fast electrical signals that cause dilation of upstream parenchymal arterioles. Stimulation of IK and SK channels within the transitional region resulted in fast-propagating conducted dilation, indicating functional expression of these channels within this vascular segment. These functional data suggest that endothelial cells that occupy the post-arteriole transitional region are phenotypically distinguishable from those found in the distal capillary tree, but a meticulous single-cell analysis is needed to verify this at the molecular level.

The NVC process is vital for maintaining cerebral blood flow to active neuronal sites. Notably, insufficient blood flow is a hallmark of many neurological disorders, including vascular cognitive impairment, stroke, and Alzheimer’s disease. The demonstration that capillaries act as sensors in the control of microcirculatory hemodynamics has altered our view of functional hyperemia (Longden et al., 2017). Our findings extend this paradigm, demonstrating that TRPA1 channels and PPADS-sensitive purinergic receptors on brain capillary endothelial cells are capable of dilating upstream parenchymal arterioles to increase RBC flux and functional hyperemia in the brain. Our findings further show that signals initiated by TRPA1 channels and purinergic receptors propagate through capillary endothelial cells through a novel Ca^2+^ signaling pathway that depends on endothelial Panx1 channels. These Ca^2+^ signals are converted to fast-conducting electrical signals by SK and IK channels in the post-arteriole transition region. We conclude that multiple sensory mechanisms present in brain endothelial cells are vital for the regulation of cerebral blood flow and functional hyperemia and could be targeted to treat cerebrovascular dysfunction.

## Materials and methods

### Chemicals and reagents

All chemicals and reagents were obtained from Sigma-Aldrich (USA) unless stated otherwise.

### Animals

Adult (12 to 16 weeks of age) male and female mice were used for all experiments. All animal procedures used in this study were approved by the Institutional Animal Care and Use Committee of the University of Nevada, Reno, School of Medicine. C57BL/6J mice (Jackson Labs) were used as wild-type controls in this study. Endothelial cell-specific deletion of TRPA1 was achieved by crossing mice homozygous for *Trpa1* containing loxP sites flanking S5/S6 transmembrane domains (Jackson Labs, stock number: 008654) with heterozygous *Tek^cre^* mice (Jackson Labs, stock number: 008863) to generate *Trpa1-*ecKO mice, as previously described (Sullivan et al., 2015). Mice homozygous for floxed *Trpa1*, but without expression of *cre*-recombinase (*Trpa1^fl/fl^*), were used as controls for experiments involving *Trpa1-*ecKO mice. Mice with tamoxifen-inducible endothelial cell-specific knockout of Panx1 channels (*Panx1*-ecKO), provided by Dr. Brant Isakson (University of Virginia, USA), were generated as previously described (Lohman et al., 2015). *Cre*-recombinase was induced by intraperitoneal (*i.p.*) injection of animals with tamoxifen (1 mg in 0.1 ml of peanut oil) on 10 consecutive days. Mice expressing *cre* alone treated with tamoxifen and *Panx1*-ecKO mice treated with vehicle were used as controls. *VEC-GCaMP8* mice, expressing the genetically encoded Ca^2+^ indicator GCaMP8 exclusively in the endothelium, were developed by CHROMus (Cornell University, USA) (Ohkura et al., 2012). Fluorescent reporter *mT/mG* mice were crossed with mice heterozygous for vascular endothelial cadherin (VEC)-*cre*, yielding mice that express EGFP in endothelial cells and tdTomato in all other cell types (Longden et al., 2017). Mice were euthanized by decapitation under isoflurane anesthesia. The brain was isolated into a solution of ice-cold, Ca^2+^-free, Mg^2+^-based physiological saline solution (Mg-PSS) containing 5 mM KCl, 140 mM NaCl, 2 mM MgCl_2_, 10 mM HEPES, and 10 mM glucose (pH 7.4, NaOH).

### Isolation of native capillary endothelial cells

Individual capillary endothelial cells were isolated as previously described (Longden et al., 2017). Brains were denuded of surface vessels with an aCSF-wetted cotton swab, and two 1-mm thick brain slices were excised and homogenized in ice-cold aCSF using a Dounce homogenizer. The brain homogenate was filtered through a 70-µM filter, and capillary networks that were captured on the filter were transferred to a new tube. Individual capillary endothelial cells were isolated by enzymatic digestion with 0.5 mg/ml neutral protease (Worthington Biochemical Corp., USA) and 0.5 mg/ml elastase (Worthington) in endothelial cell isolation solution composed of 5.6 mM KCl, 55 mM NaCl, 80 mM sodium glutamate, 2 mM MgCl_2_, 0.1 mM CaCl_2_, 4 mM glucose, and 10 mM HEPES (pH 7.3) for 45 minutes at 37°C. After the first digestion, 0.5 mg/ml type I collagenase (Worthington) was added, and a second 2-minute incubation at 37°C was performed. Digested networks were washed in ice-cold endothelial cell isolation solution, then triturated with a fire-polished glass Pasteur pipette to produce individual endothelial cells.

### Whole-cell patch-clamp electrophysiology

Whole-cell patch-clamp electrophysiology was used to assess the presence of functional TRPA1 channels on native brain capillary endothelial cells. Following isolation, capillary endothelial cells were transferred to a recording chamber and allowed to adhere to glass coverslips for 10 minutes at room temperature (∼22°C).

TRPA1 currents were recorded in cells patch-clamped in the conventional whole-cell configuration, and currents were amplified using an Axopatch 200B amplifier. Currents were filtered at 1 kHz and digitized at 10 kHz. Pipettes were fabricated from borosilicate glass (1.5 mm outer diameter, 1.17 mm inner diameter; Sutter Instruments, USA), fire-polished to yield a tip resistance of 3-6 MΩ and filled with a solution consisting of 10 mM NaOH, 11.4 mM KOH, 128.6 mM KCl, 1.091 mM MgCl_2_, 3.206 mM CaCl_2_, 5 mM EGTA, 10 mM HEPES and 5 mM ruthenium red (pH 7.2). The bath solution consisted of 134 mM NaCl, 6 mM KCl, 2 mM MgCl_2_, 10 mM HEPES, 4 mM glucose, and 1 mM EGTA (pH 7.4). Currents were induced by a ramp protocol (−100 to +100 mV in 10-mV steps, 300 ms per step). The mean capacitance of capillary endothelial cells was 6.83 ± 2.2 pF (n = 7 cells from 3 animals). TRPA1 currents were recorded following the addition of 4-HNE (100 nM; Cayman Chemical, USA) to the external bath solution. The selective TRPA1 antagonist HC-030031 (10 µM; Tocris, USA) was used to validate the current.

IK, SK, and K_ir_ *c*urrents were recorded using the conventional whole-cell configuration at a holding potential of −50 mV, with 400-ms ramps from −100 to +45 mV. The external bathing solution was composed of 134 mM NaCl, 6 mM KCl, 1 mM MgCl_2_, 2 mM CaCl_2_, 10 mM glucose, and 10 mM HEPES. The composition of the pipette solution was 10 mM NaCl, 30 mM KCl, 10 mM HEPES, 110 mM K^+^ aspartate and 1 mM MgCl_2_ (pH 7.2). IK and SK currents were activated by adding NS309 (10 µM) to the external bath solution. K_ir_ channel currents were recorded with elevated extracellular [K^+^] in the external bath solution (60 mM KCl, 80 mM NaCl). BaCl_2_ (10 µM) was used to isolate K_ir_ currents.

Clampex and Clampfit (version 10.2; Molecular Devices) were used for data acquisition and analysis, respectively. All recordings were performed at room temperature.

### Pressure myography

Pressure myography was performed using the recently described arteriole-capillary microvascular preparation (Longden et al., 2017), consisting of a capillary segment with attached, intact capillaries (Figure 2A). Briefly, a 3 mm x 5 mm x 3 mm (W x L x D) section of the brain surrounding the middle cerebral artery was dissected and placed into a microdissection dish containing ice-cold Mg-PSS. The middle cerebral artery and the surrounding pial meninge were carefully removed so as to maintain branching parenchymal arterioles within the brain tissue. Thereafter, parenchymal arterioles with attached capillaries were carefully blunt-dissected from the underlying cerebral tissue and transferred to a pressure myograph chamber containing oxygenated (21% O_2_, 6% CO_2_, 73% N_2_) aCSF (124 mM NaCl, 3 mM KCl, 2 mM CaCl_2_, 2 mM MgCl_2_, 1.25 mM NaH_2_PO_4_, 26 mM NaHCO_3_, 4 mM glucose) and a Sylgard pad at the chamber base. Microvascular preparations were mounted between two glass micropipettes (outer diameter, ∼20-40 μm) and secured with nylon monofilaments. The outermost tips of attached capillaries were pinned onto the surface of the Sylgard pad to immobilize the capillary bed and allow pressurization of the preparation. Intraluminal pressure was controlled using a servo-controlled peristaltic pump (Living Systems Instrumentation, USA). Pressurized arteries were visualized with an inverted microscope (Accu-Scope Inc., USA) coupled to a USB camera (The Imaging Source LLC, USA). Intraluminal diameter as a function of time was recorded using IonWizard software (version 6.4.1.73; IonOptix LLC, USA). Microvascular preparations were bathed in warmed (37°C), oxygenated aCSF at an intraluminal pressure of 5 mmHg. Following a 15-minute equilibration period, intraluminal pressure was increased to 40 mmHg and spontaneous tone was allowed to develop. Preparations that developed less than 10% tone were discarded. Localized application of drugs onto the capillary bed was achieved by placing a micropipette attached to a Picospritzer III (Parker, USA) adjacent to capillary extremities. The viability of attached capillaries was assessed by locally applying a 7-second pulse of aCSF containing elevated [K^+^] (10 mM) onto capillary extremities. Preparations that failed to dilate to elevated [K^+^] were discarded.

In control experiments designed to validate the microvascular preparation, the connection between the upstream parenchymal arteriole segment and attached capillaries was severed. In other control experiments, spatial spread was determined by pulsing aCSF containing 1% w/v Evans Blue dye. The role of TRPA1 channels was examined by locally applying 4-HNE (1 µM) or AITC (30 µM) to the attached capillaries, and the role of purinergic signaling was examined by focal application of ATP (10 µM). Underlying mechanisms were examined by adding pharmacological agents to the superfusing bath solution. In separate experiments, preparations were modified by removing attached capillary segments and leaving the post-arteriole transitional segment intact. In these experiments, the post-arteriole transitional segment was directly stimulated via a micropipette placed adjacent to this segment. Lumen diameter was continuously recorded, and responses were expressed as vasodilation (%). Velocity data were obtained by determining the latency of the response following local application of compounds in relation to the distance traveled between the capillary extremity or post-arteriole transitional segment to the parenchymal arteriole.

### Fluorescence imaging

Images of the *ex vivo* microvascular preparation were obtained from VEC-*cre mT/mG* mice. Images were obtained using a custom-built upright microscope (Olympus BX51 WI; Olympus Corp., Japan) equipped with epifluorescence illumination (CoolLED pE-300^white^, CoolLED Ltd., UK), and an ORCA-Fusion Digital CMOS C14440-20UP camera (Hamamatsu Corporation, Japan). Images of the complete microvascular preparation were obtained using a 20x water immersion objective (numerical aperture 0.5, Olympus), and magnified images of the different vascular segments using a 60x water immersion objective (numerical aperture 1.0, Olympus). Each field of view was 1152 x 1152 pixels (20x, 1 pixel = 0.65 μm; 60x, 1 pixel = 0.22 μm). Images were captured using μManager software (version 1.4.22, University of California, San Francisco, USA) and analyzed using ImageJ software (version 1.52n, National Institutes of Health, USA).

The length of capillary endothelial cells was determined by fluorescence labeling isolated capillary networks. Isolated capillary networks were placed onto a glass coverslip and fixed (4% formaldehyde in phosphate buffered saline (PBS)) and blocked (0.5% Triton X-100, 5% SEABlock Blocking Buffer (ThermoFisher Scientific, USA) in PBS, 1 hour on ice). Capillary networks were visualized by staining preparations with Alexa Fluor^TM^ 568 conjugated isolectin B4 (1:200 (catalogue number I21412, ThermoFisher Scientific), 2 hours at room temperature). Aqueous mounting medium with DAPI (Abcam, USA) was used to visualize nuclei of endothelial cells. Fluorescence images were taken by an Olympus FluoView laser scanning confocal microscope (Olympus FluoView FV1000, Olympus) using a 60x oil-immersion objective (numerical aperture: 1.42, Olympus). Each field of view was 1024 x 1024 pixels (1 pixel = 0.21 μm). Z-stacks were captured at 50 µm thickness to allow for 3-dimensional reconstruction of the capillary networks. Images were captured using FluoView FV1000 FV10-ASW software (version 4.02, Olympus), and analyzed using ImageJ software. The length of capillary endothelial cells was determined by measuring the distance between nuclei.

### Ca2+ imaging

Microvascular preparations isolated from *VEC-GCaMP8* mice were mounted in a pressure myograph chamber, as described above. Changes in capillary endothelial cell [Ca^2+^] were recorded in real time after local exposure to TRPA1 agonists or ATP. Movies were recorded at an average of 23 frames/s for approximately 120 seconds (2760 frames). Baseline Ca^2+^ activity was recorded for the first 60 seconds (∼1380 frames), then the capillary bed was picospritzed with AITC (30 µM) or ATP (10 µM), and the preparations were recorded for an additional 60 seconds (∼1380 frames). Movies were obtained using a custom-built upright microscope (Olympus BX51 WI) equipped with epifluorescence illumination (CoolLED pE-300^white^), a 20x water immersion objective (numerical aperture 0.5, Olympus), and an ORCA-Fusion Digital CMOS C14440-20UP camera. Each field of view was 1152 x 1152 pixels (1 pixel = 0.65 μm). Movies were captured using μManager software (version 1.4.22, University of California, San Francisco, USA) and analyzed using ImageJ software (version 1.52n, National Institutes of Health, USA). Discrete regions of interest (ROIs) along the capillary network were selected for analysis. The fractional increase in fluorescence (F/F_0_) was determined for ROIs, where fluorescence (F) is normalized to basal fluorescence (F_0_). Underlying mechanisms were examined by adding pharmacological agents to the superfusing bath solution. Velocity data were obtained by determining the latency of the response between adjacent ROIs following local application of compounds in relation to the distance traveled.

### Two-photon imaging of in vivo brain microcirculatory hemodynamics

Hemodynamics of the murine microcirculation were assessed *in vivo* after capillary endothelial cell TRPA1 activation as described previously (Longden et al., 2017). Briefly, mice were anesthetized with isoflurane (5% induction, 2% maintenance) and the skull was exposed. Thereafter, a stainless-steel head plate was attached over the right hemisphere using dental adhesive, and the head was immobilized by securing the head plate to a holding frame. A cranial window (∼2 mm diameter) was made in the skull above the somatosensory cortex, after which FITC-labeled dextran (150 kDa; 150 μL of a 3 mg/mL solution) was intravenously (*i.v.*) administered to allow visualization of the cerebral vasculature and contrast imaging of RBCs. Following completion of the surgical procedure, isoflurane anesthesia was replaced with combined α-chloralose (50 mg/kg, *i.p.*) and urethane (750 mg/kg, *i.p.*) to eliminate confounding vasodilatory effects of isoflurane. Capillaries downstream of arterioles were identified and selected for study. The duration and pressure of fluid ejection via a glass micropipette was calibrated to obtain a localized application area approximately 10 μm in diameter. The micropipette was positioned adjacent to the capillary, and AITC (30 µM) was directly applied onto the capillary. Tetramethylrhodamine isothiocyanate (TRITC)-labeled dextran was included in the micropipette to determine the spatial coverage of the solution. Images were obtained using a Zeiss LSM-7 two-photon microscope (Zeiss, USA), equipped with a 20x Plan Apochromat 1.0 NA DIC VIS-IR water-immersion objective (Zeiss) and coupled to a Coherent Chameleon Vision II Titanium-Sapphire pulsed infrared laser (Coherent, USA). After excitation at 820 nm, emitted FITC-labeled dextran and TRITC-labeled dextran fluorescence was separated through 500-550 and 570-610 nm bandpass filters, respectively. RBC flux was determined by line-scan imaging of the capillary and diameter changes of upstream arterioles were obtained.

### Functional hyperemia

After anesthetizing mice with isoflurane (5% induction, 2% maintenance), the skull was exposed and the head was immobilized in a stereotaxic frame. The skull of the right hemisphere was carefully thinned using a drill to visualize the surface vasculature of the somatosensory cortex. Following completion of the surgical procedure, isoflurane anesthesia was replaced with combined α-chloralose (50 mg/kg, *i.p.*) and urethane (750 mg/kg, *i.p.*) to eliminate confounding vasodilatory effects of isoflurane. Perfusion was monitored via a laser-Doppler flowmetry probe (PeriFlux System PF5000, Perimed AB, Sweden) positioned above the somatosensory cortex. The contralateral whiskers were stimulated for ∼20 seconds, and changes in perfusion were recorded. Contralateral whiskers were stimulated three times at 2-minute intervals. Ipsilateral whiskers were also stimulated as a control for potential vibration artifacts. The role of TRPA1 channels was assessed by treating mice with HC-030031 (100 mg/kg, *i.p.* for 30 minutes). Data are presented as changes in perfusion relative to baseline, calculated as follows: %Δ Blood flow = (average perfusion during stimulus/average baseline perfusion) × 100.

### Statistical analysis

All data are expressed as means ± standard error of the mean (SEM), unless specified otherwise. The value of “n” refers to number of cells for patch-clamp electrophysiology experiments, the number of vessel preparations for pressure myography and Ca^2+^ experiments, and the number of animals for two-photon imaging studies and functional hyperemia assessments. Statistical analyses and graphical presentations were performed using GraphPad Prism v8.2 (GraphPad Software, Inc., USA). Statistical analyses were performed using Students *t*-test or analysis of variance (ANOVA), and a value of P < 0.05 was considered statistically significant.

## Authors contributions

S.E. initiated and supervised the project. S.A. and H.A.T.P. performed patch-clamp electrophysiology experiments. P.T. and P.W.P. performed pressure myograph experiments. P.T. and M.G.A. performed fluorescence and Ca^2+^ imaging experiments. A.M. performed *in vivo* two-photon laser-scanning microscopy experiments. P.T. and C.H.T. performed *in vivo* functional hyperemia experiments. B.E.I. provided *Panx1*-ecKO mice. P.T. and S.E. wrote the manuscript and prepared the figures. P.T. and S.E. revised the manuscript.

