## Supplemental Figure for "Brain Endothelial Cell TRPA1 Channels Initiate Neurovascular Coupling"

**This PDF file includes:**

Supplemental Figures 1 to 8
Captions for Supplemental Movies 1 to 10

**Other Supplementary Materials for this manuscript include the following:**

Supplemental Movies 1 to 10

### Supplemental Figures

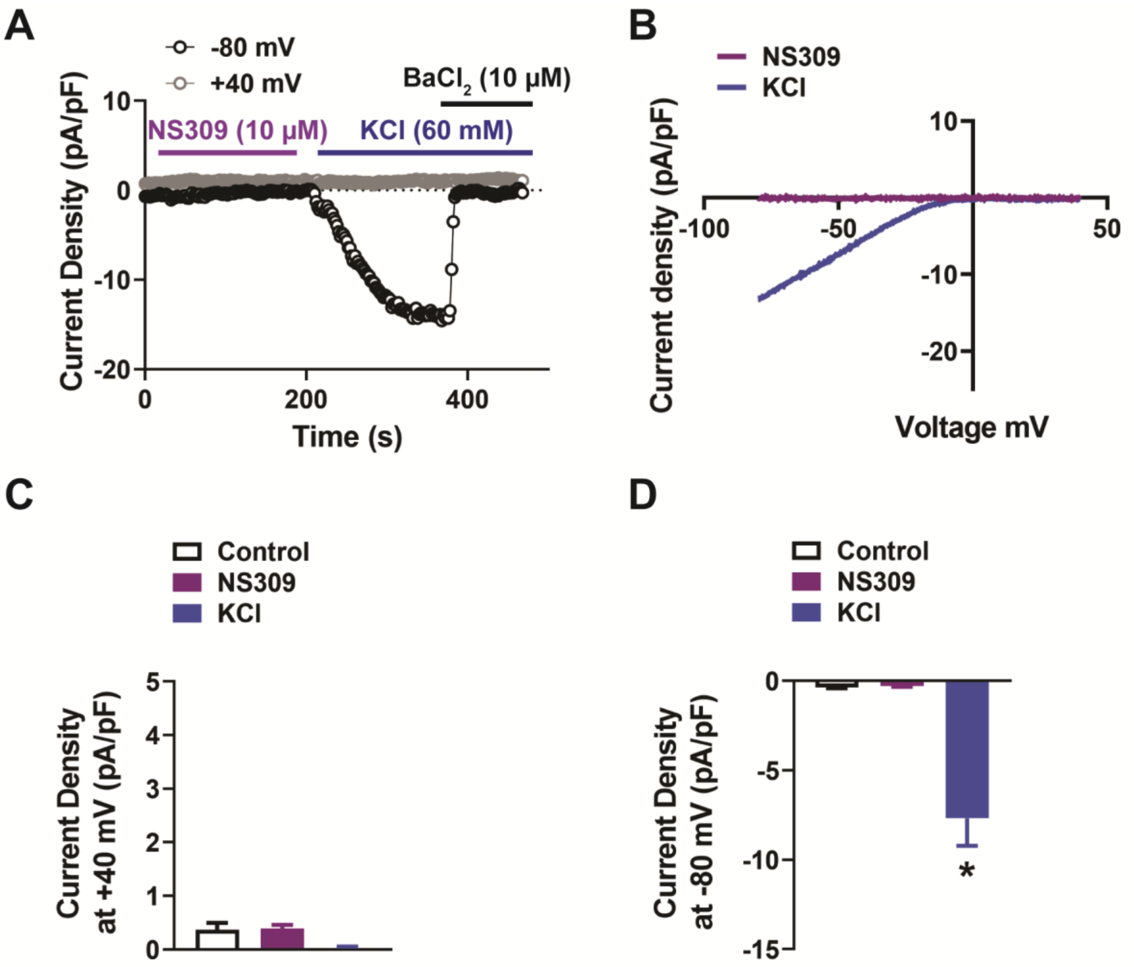


**Supplemental Figure 1. Lack of functional IK and SK channels in native capillary endothelial cells.** A and B) Representative current-time trace (A) and *I-V* relationship (B) from a whole-cell patch-clamp electrophysiology experiment demonstrating that the IK and SK channel activator NS309 (10 µM) did not produce a current in isolated native capillary endothelial cells from wild-type mice. $K_{ir}$ currents were detected by raising extracellular KCl (60 mM) and were confirmed by assessing their sensitivity to $BaCl_2$ (10 µM). C and D) Summary data showing the current produced by NS309 (10 µM) and KCl (60 mM) at +40 mV (C) and -80 mV (D) (n = 6 cells from 3 animals; *P < 0.05).

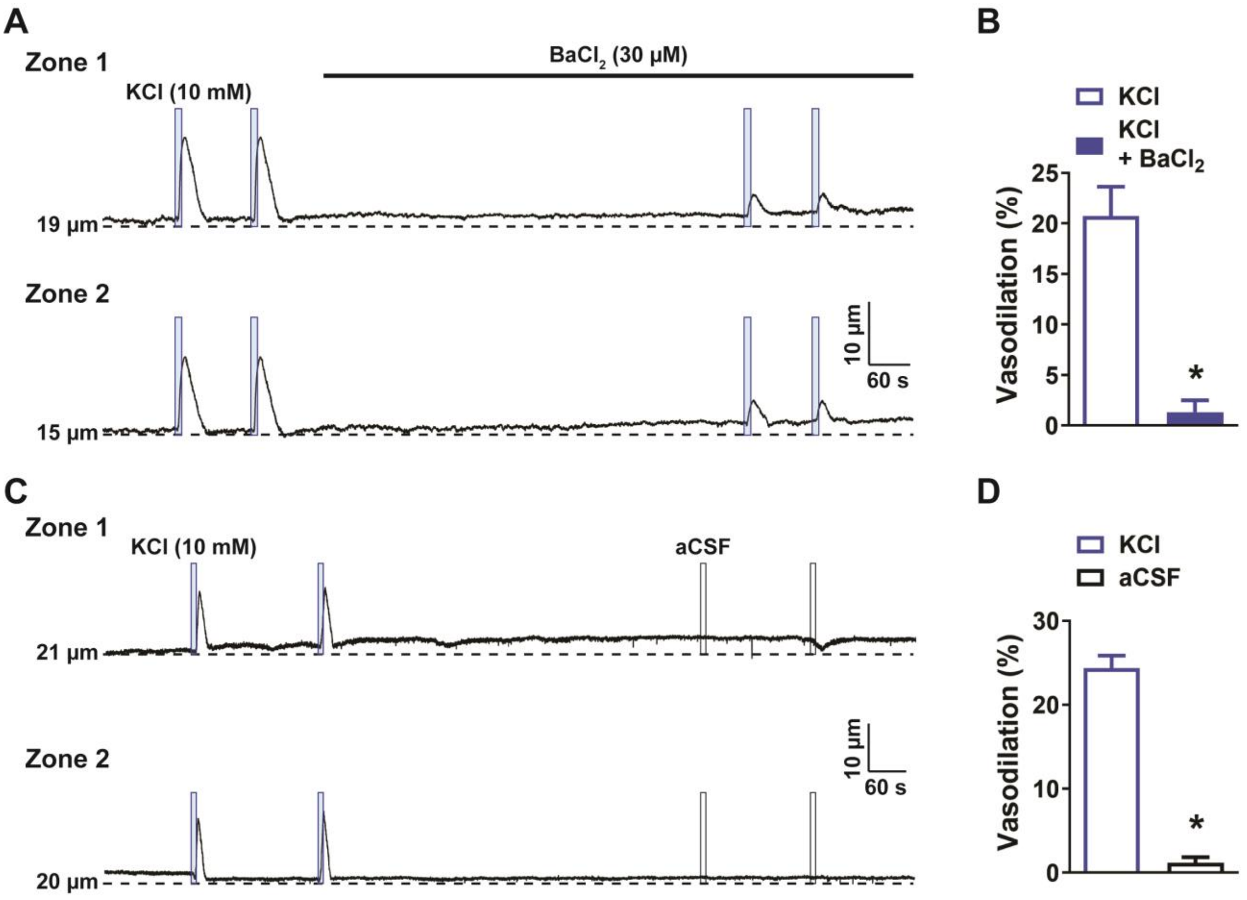


**Supplemental Figure 2. KCl-induced dilation of upstream arterioles is blocked by $BaCl_2$.** A) Representative traces (A) and summary data (B) showing that application of a solution containing elevated KCl (10 mM; blue box) directly onto capillary extremities induced a dilation of the upstream arteriole that was blocked by $BaCl_2$ (30 μM) in *ex vivo* microvascular preparations from wild-type mice (n = 6 preparations from 4 animals; *P < 0.05). C and D) Representative trace (C) and summary data (D) showing that application of aCSF (black box) onto the capillary bed did not affect the lumen diameter of upstream arterioles (n = 6 preparations from 4 animals; *P < 0.05).

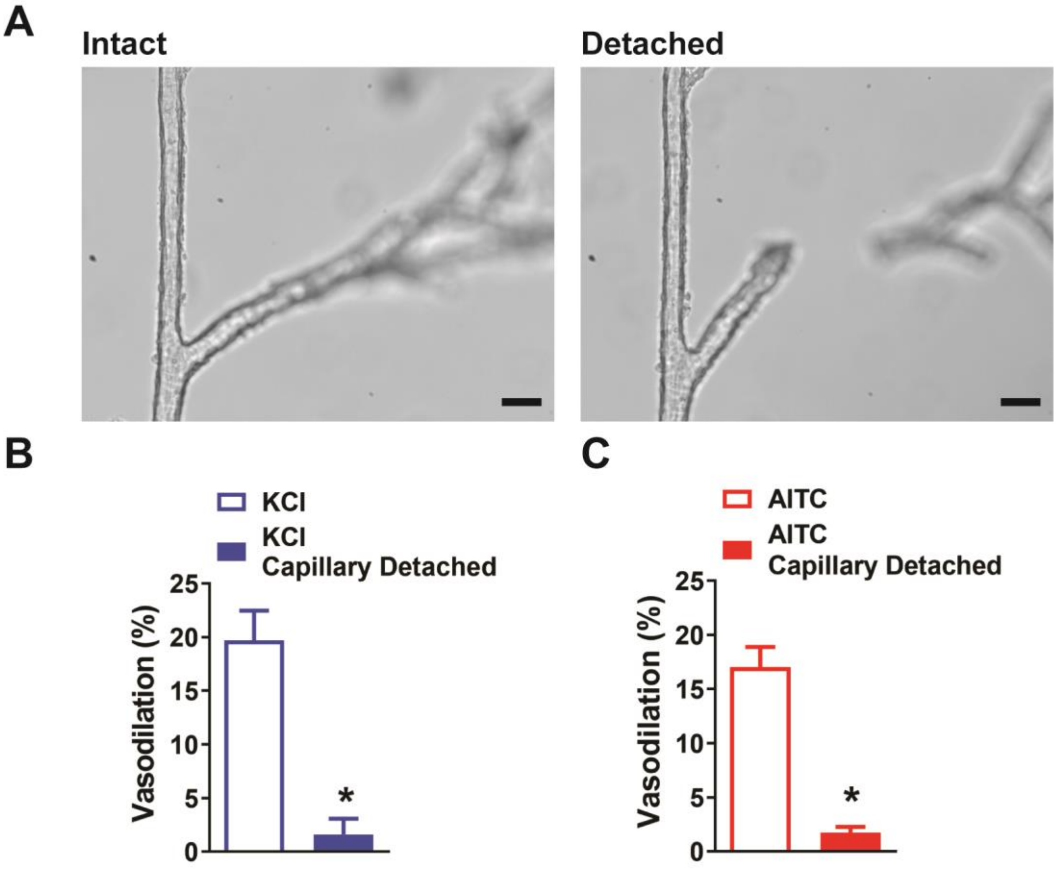


**Supplemental Figure 3. Severing the connection between capillaries and the arteriole segment of an ex vivo microvascular preparation.** A) Representative images showing an intact microvascular preparation (*left*) and the same preparation after severing the connection between the arteriole and capillary (*right*). Scale bar = 50 µm. B and C) Summary data demonstrating the loss of propagative dilation to elevated KCl (10 mM) (n = 7 preparations from 5 animals; *P < 0.05) (B) and AITC (30 µM) (n = 7 preparations from 5 animals; *P < 0.05) (C) after the connection is lost.

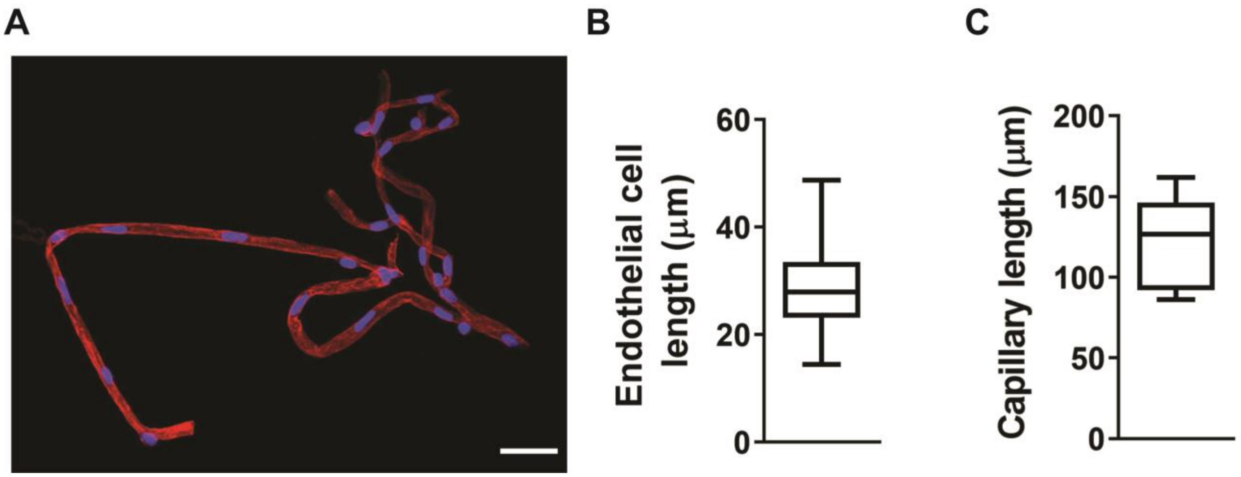


**Supplemental Figure 4. Average length of a cerebral capillary endothelial cell.** A) Representative image of an isolated cerebral capillary network. Endothelial cells and nuclei were stained with isolectin B4 (red) and DAPI (blue), respectively. Scale bar = 25 μm. B) Summary data showing the average length of capillary endothelial cells, calculated by measuring the inter-nuclear distance (determined from 173 endothelial cells from 4 animals). C) Summary data showing the average length of the capillary segment in isolated *ex vivo* microvascular preparations (n = 9 preparations from 6 animals). Data are shown as a box and whisker plot.

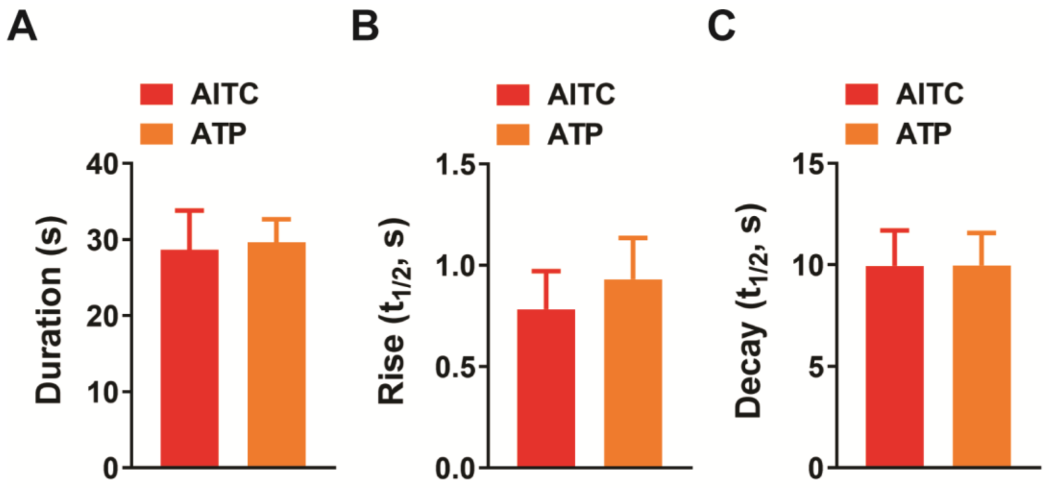


**Supplemental Figure 5. Kinetics of the $Ca^{2+}$ response in capillaries following activation of TRPA1 channels purinergic receptors.** A to C) Duration (A), rise time [half-time ($t_{1/2}$, s)] (B) and decay time ($t_{1/2}$, s) (C) of the $Ca^{2+}$ response following focal application of AITC (30 µM) and ATP (10 µM) to distal capillaries in microvascular preparations from *VEC-GCaMP8* (AITC, n = 6 preparations from 5 animals; ATP, n = 13 preparations from 12 animals).

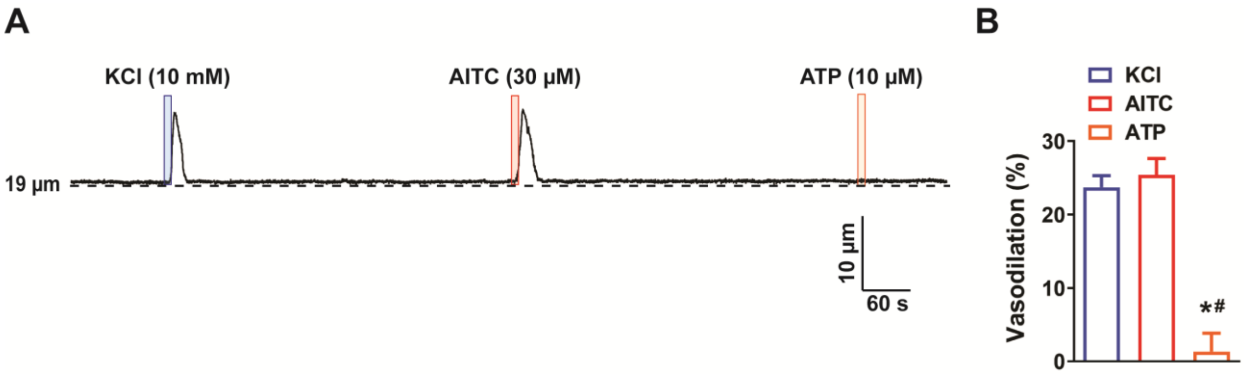


**Supplemental Figure 6. ATP does not evoke dilation of upstream arterioles when applied to the post-arteriole transitional segment.** A) Representative trace showing that application of elevated KCl (10 nM; blue box) or AITC (30 μM; red box) onto the post-arteriole transitional segment increased the lumen diameter of the upstream arteriole, but application of ATP (10 μM; orange box) to this segment did not. B) Summary data showing that application of ATP (10 μM) onto the post-arteriole transitional segment did not dilate upstream arterioles (n = 6 preparations from 3 animals; *P < 0.05 vs. KCl, #P < 0.05 vs. AITC).

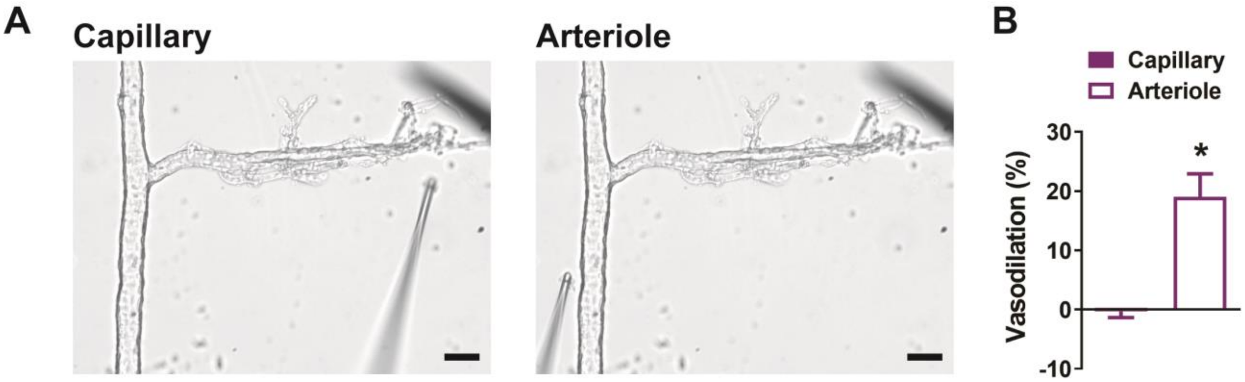


**Supplemental Figure 7. Application of NS309 to capillary extremities has no effect on the upstream arteriole.** A) Representative images of an intact microvascular preparation showing the drug-administering cannula positioned adjacent to capillary extremities (*left*) and arteriole segment (*right*). Scale bar = 50 µm. B) Summary data showing that application of the IK and SK channel activator NS309 (10 µM) did not induce relaxation of the upstream arteriole when applied to capillaries, but did cause dilation when applied directly to the arteriole (n = 6 preparations from 3 animals; *P < 0.05).

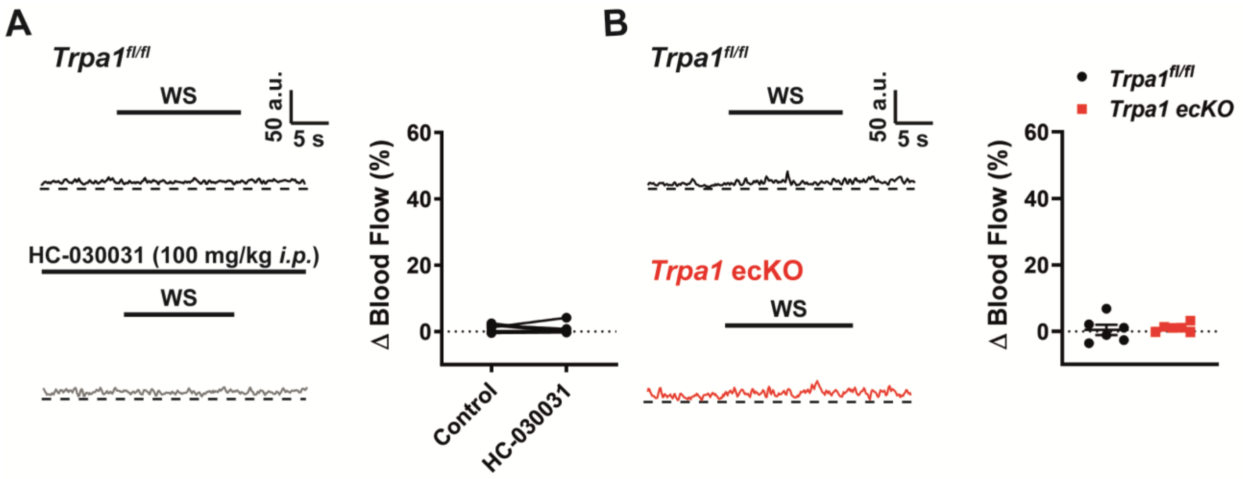


**Supplemental Figure 8. Lack of a hyperemic response following ipsilateral whisker stimulation.** A) Representative traces (*left*) and summary data (*right*) showing the lack of a hyperemic response in the somatosensory cortex following ipsilateral whisker stimulation (WS), measured using laser-Doppler flowmetry in mice prior to and following treatment with HC-030031 (100 mg/kg *i.p.*) (n = 6 animals). B) Representative traces (*left*) and summary data (*right*) showing the lack of a hyperemic response in the somatosensory cortex following ipsilateral whisker stimulation (WS), measured using laser-Doppler flowmetry in control *Trpa1*$^{fl/fl}$ mice and *Trpa1*-ecKO mice (n = 6 animals).

### Supplemental movie legends

**Supplemental Movie 1. Localized application of Evans Blue dye onto capillary extremities of a microvascular preparation.** Representative time-series images of a microvascular preparation demonstrating that application of Evans Blue dye (1% w/v) is localized to the region of the capillary tree and does not spread to the upstream parenchymal arteriole. Evans Blue was applied at 10 seconds. Scale bar = 100 µm.

**Supplemental Movie 2. Localized application of AITC onto capillary extremities of a microvascular preparation dilates the upstream arteriole.** Representative time-series images of a microvascular preparation demonstrating that localized application of AITC (30 µM) onto capillary extremities dilates the upstream arteriole. AITC was applied at 10 seconds. Scale bar = 50 µm.

**Supplemental Movie 3. Focal application of AITC onto capillary extremities induces an increase in intracellular [$Ca^{2+}$].** Representative time-series images of a microvascular preparation demonstrating that localized application of AITC (30 µM) onto distal capillary extremities produces a propagative $Ca^{2+}$ signal. AITC was applied at 10 seconds. Scale bar = 50 µm.

**Supplemental Movie 4. Increase in intracellular [$Ca^{2+}$] initiated by AITC is blocked by the selective TRPA1 antagonist HC-030031.** Representative time-series images of a microvascular preparation demonstrating that the propagative $Ca^{2+}$ signal produced by AITC (30 µM) is blocked by suerpfusing the preparation with HC-030031 (10 µM). AITC was applied at 10 seconds. Scale bar = 50 µm.

**Supplemental Movie 5. Focal application of ATP onto capillary extremities induces an increase in intracellular [$Ca^{2+}$].** Representative time-series images of a microvascular preparation demonstrating that localized application of ATP (10 µM) onto distal capillary extremities produces a propagative $Ca^{2+}$ signal. ATP was applied at 10 seconds. Scale bar = 50 µm.

**Supplemental Movie 6. Increase in intracellular [$Ca^{2+}$] initiated by AITC is blocked by the pan-P2X inhibitor PPADS.** Representative time-series images of a microvascular preparation demonstrating that the propagative $Ca^{2+}$ signal produced by ATP (10 µM) is blocked by superfusing the preparation with PPADS (10 µM). ATP was applied at 10 seconds. Scale bar = 50 µm.

**Supplemental Movie 7. ATP-induced $Ca^{2+}$ signal under normal conditions.** Representative time-series images of a microvascular preparation demonstrating that localized application of ATP (10 µM) onto distal capillary extremities produces a propagative $Ca^{2+}$ signal in preparations superfused with $Ca^{2+}$-containing aCSF. ATP was applied at 10 seconds. Scale bar = 50 µm.

**Supplemental Movie 8. ATP-induced $Ca^{2+}$ signal is abolished in extracellular $Ca^{2+}$-free conditions.** Representative time-series images of a microvascular preparation demonstrating that localized application of ATP (10 µM) onto distal capillary extremities failed to induced a propagative $Ca^{2+}$ signal in preparations superfused with $Ca^{2+}$-free aCSF. ATP was applied at 10 seconds. Scale bar = 50 µm.

**Supplemental Movie 9. ATP-induced $Ca^{2+}$ signal is restored following reintroduction of extracellular $Ca^{2+}$.** Representative time-series images of a microvascular preparation demonstrating that the propagative $Ca^{2+}$ signal following localized application of ATP (10 µM) onto distal capillary extremities returned once extracellular $Ca^{2+}$ (2 mM) was reintroduced to the preparation. ATP was applied at 10 seconds. Scale bar = 50 µm.

**Supplemental Movie 10. Localized application of Evans Blue dye onto the post-arteriole transitional segment of a microvascular preparation.** Representative time-series images of a modified microvascular preparation in which the capillary tree was removed. Application of Evans Blue dye (1% w/v) onto the post-arteriole transitional segment was localized to this region and did not spread to the upstream parenchymal arteriole. Evans Blue was applied at 10 seconds. Scale bar = 100 µm.
